# Point cloud local ancestry inference (PCLAI): continuous coordinate-based ancestry along the genome

**DOI:** 10.64898/2026.03.23.713813

**Authors:** Margarita Geleta, Daniel Mas Montserrat, Nilah M. Ioannidis, Alexander G. Ioannidis

## Abstract

Local ancestry inference (LAI) predicts a discrete ancestry label for each segment of an individual’s genome and has become integral to studying population history, genetic variation, and polygenic trait association. We present a new local ancestry paradigm that eschews discrete categorical labels and instead performs inference in a continuous coordinate space. We call this method “point cloud local ancestry inference” (PCLAI), since it represents an individual’s genetic ancestry as a point cloud with each point corresponding to a small haplotypic segment in their genome. This formulation works in any co-ordinate space (for instance, geographic or principal components) permitting the representation of continuous genetic variation at the haplotypic-segment level without resorting to artificially constructed discrete labels. We illustrate PCLAI by training on ancient samples from multiple time periods separately, yielding chromosome paintings based on geography that are time-stratified and provide insight into how individuals’ genomic segments moved across space and time.

## Introduction

Advancements in genetics have shown that human origins are more complex than previously thought. Rather than simple branching lineages consistent with early formulations of the out-of-Africa model, multiple lines of evidence now reveal a reticulate evolutionary history with extensive genetic interaction between lineages [1, 2, 3]. Following admixture between populations, recombination reshapes individual genomes into mosaics of haplotypic segments inherited from different ancestral sources. Shared chromosomal segments that are identical-by-descent connect individuals to recent relatives, with expected segment lengths shrinking as the number of recombination events separating two haplotypes increases; consequently, more distant genealogical connections are increasingly represented by short segments that are harder to reliably detect and interpret [4, 5].

In population genetics, ancestry has often been framed using population descriptors as proxies for genetic similarity to investigator-constructed reference categories [6], often selected based on differentiation through isolation, founder effects, and endogamy. These factors shape allele frequencies and haplotype linkage structure and thereby influence downstream inference, including applications in precision medicine where differences between genetic ancestries have been shown to be significant in drug response and polygenic risk score accuracies [7, 8, 9]. However, in many cases investigator-driven ancestry classification schemes are influenced by socially constructed ethnic ontologies, rather than genetic clustering, thus mixing non-genetic factors and genetic factors into the definition of genetic ancestry covariates.

When considering genetic ancestry, two types are commonly encountered: global and local. Global ancestry inference methods assign individuals to a single point, whether in a principal component analysis (PCA) space, a UMAP space [10], or a clustering space defined by a vector of mixture proportions. For clustering, the Bayesian MCMC clustering *STRUCTURE* [11] and its maximum-likelihood analogue *ADMIXTURE* [12] fit closely related latent-variable models but differ substantially in scalability, with *ADMIXTURE* optimized for fast analysis of large cohorts. More recent accelerations such as *Neural ADMIXTURE* [13] or *ADAMIXTURE* [14] recast the *ADMIXTURE* objective in an autoencoder formulation, obtaining similar cluster assignments with substantially reduced runtime, while *HaploADMIXTURE* [15] incorporates short-range haplotypic information to better leverage linkage disequilibrium (LD) instead of treating markers as independent. Other neural network approaches include the fully-connected *Diet Networks* [16] and *ægen* [17], which uses variational autoencoders to infer global ancestry labels. These methods detect allele frequency differences within the data and assign an individual’s whole genome to a set of subpopulation proportions.

In parallel, the work of Cavalli-Sforza [18] and Novembre [19] demonstrated that genetic variation exhibits a strikingly continuous component: principal component analysis (PCA) decomposition of genetic variation captures population structure that often recapitulates broad geographic patterns. Indeed, spatial methods can explicitly model allele frequencies as smooth functions over continental geography, as in spatial ancestry analysis *SPA* [20]. More recently *Locator* [21] showed that deep neural networks can predict geographic coordinates directly from genotypes, but the method struggles with intercontinental admixture, since an individual with multiple ancestries across their genome is plotted as an average of those ancestries, sometimes falling in implausible intermediate locations like the middle of an ocean. In our work, we argue that the geographic-genetic correlation is not generalizable across all geographies and populations, although it can hold under specific isolation-by-distance scenarios. Furthermore, ancestry is a dynamic and continuously evolving descriptor: the geographic-genetic associations of populations of the past are very different from those today [3, 22], a fact that reinforces that population descriptors not only vary spatially, but also temporally—both continuous dimensions for which the commonly accepted discretization of genetic ancestry is a lossy representation of its richness and variability [23].

Human genomes present a mosaic structure produced by recombination, which global ancestry methods discussed above ignore, as they assign each individual only a genome-wide average ancestry descriptor. Local ancestry inference (LAI) unravels this phenomenon by partitioning the genome into segments with each segment then assigned separately to discrete ancestry templates, typically formulated as a sequence labeling problem with discrete ancestry labels. Early LAI methods largely used generative, Markovian models of ancestry transitions along the genome—e.g., the hidden Markov model (HMM) in *SABER* [24] to account for ancestral LD, and related HMM-based approaches such as *HAPAA* [25], *HAPMIX* [26], *PCAdmix* [27], and the more recent *FLARE* [28] that model local ancestry as latent states with transitions driven by recombination. Window-based strategies such as *LAMP* [29] trade some modeling fidelity for speed by operating on local genotype and haplotype summaries. Later discriminative methods improved robustness and accuracy by learning direct mappings from haplotypes to labels, notably Durand’s SVM-based approach [30] and tree-based *RFMix* [31] and *Gnomix* [32]. There is also *Loter* [33] based on an optimization problem solved via dynamic programming. Deep learning approaches were introduced via convolutional models in *LAI-Net* [34], the first neural network to explore LAI, the template-matching network *SALAI-Net* [35], which showed transferability across populations and species, and the transformer-based *Orchestra* [36], which extended ideas explored in SALAI-Net, further improving accuracy and throughput for biobank-scale cohorts. Despite their differences, all these methods share their fundamental conception of the problem, viewing ancestry as a discrete label drawn from a set of investigator-defined reference categories.

These strands suggest a reframing: admixed genomes can be modeled as sequences of discrete recombination events (breakpoints) joining segments whose local descriptors should vary continuously in a genetic embedding space. In this view, each continuous segment along the genome can be represented by a coordinate vector in a low-dimensional space, and an individual’s ancestry descriptor becomes a *point cloud* of coordinates indexed along their genome, separated by discrete recombination breakpoints. We therefore propose *point cloud local ancestry inference* (*PCLAI*), where, rather than predicting a discrete ancestry label per window, we regress a coordinate for each single nucleotide polymorphism (SNP) window in a continuous space while simultaneously detecting breakpoints that mark recombination events. Conceptually, each predicted point is a local genomic coordinate and each breakpoint is a discrete junction—together yielding a representation that is both continuous within haplotypic blocks and discrete at boundaries. To make such a high resolution continuous local ancestry model tractable and scalable, we follow a window-based approach leveraging transformer representations to assign spatially continuous ancestry descriptors to every genomic position. We regress PCA coordinates per chromosome window, with an *L*_1_ objective on coordinates and a permutation-invariant Chamfer dissimilarity term [37] that aligns predicted and target point clouds in the embedding space. This design directly mitigates the problem of global PCA projections, which cease to visibly encode admixture after modest numbers of generations, even though the local mosaic structure persists [5]; PCLAI is explicitly built to preserve that locus-specific information. This framing integrates the interpretability of coordinate-based structure with the discrete resolution of recombination events, providing a unified substrate for analyzing admixture, and representing how ancestry coordinates vary across both space and time.

## Methods

### PCLAI model overview

PCLAI can paint an individual’s chromosomes in any target vector space, assigning each haplotypic segment a coordinate vector that could derive from a physical (geographic) space defined by latitude and longitude or an abstract, genetic ancestry related coordinate space, such as PCA or UMAP. To make PCLAI tractable and scalable on whole-genome SNP array data, we represent each phased haplotype strand 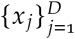 (with *D* SNPs) as a sequence of fixed-length, non-overlapping windows 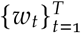 with *T* = ⌈*D* / *h*⌉, where *h* is the number of SNPs per window (we use *h* = 500 and 1,000 in our experiments), so a window *w*_*t*_ ∈ ℝ^*h*^ contains the SNP subsequence *x* (_*t*− 1_) _*h* + 1_, …, *x*_min_ (_*t* · *h,D*_), zero-padded on the right if *t* · *h* > *D*. Because SNP density varies along the genome, a fixed-*h* window corresponds to a variable number of base pairs; we choose *h* so that windows typically span on the order of 100 kb (∼ 10^5^ base pairs). The raw diploid genotypes are first phased and split into haplotype strands, after which each haplotype is processed independently. Working with windows is beneficial because it captures local structure such as LD. The non-overlapping, windowed representation serves two purposes: it reduces sequence length and attention cost while preserving local haplotypic context, and it provides a natural unit at which to predict both: a continuous ancestry coordinate and a breakpoint probability.

For each haplotypic strand, each SNP window *w*_*t*_ ∈ ℝ^*h*^ is passed through a shared fully-connected encoder (**Figure 1.A**). The first layer maps the *h*-dimensional window into a hidden representation using rectified linear units and layer normalization [38] to stabilize training, and the second projects to the transformer model dimension *d*_model_. This produces, for each haplotype, a sequence of token embeddings, one per window. The encoder shares weights across windows and across haplotypes. Because the number of windows per chromosome can vary, the model operates on padded sequences. For each batch, we construct binary masks indicating which positions correspond to real windows versus padding; these masks are propagated through the attention mechanism and applied in the breakpoint head so that padding does not contribute to the attention scores or the loss.

**Figure 1.**
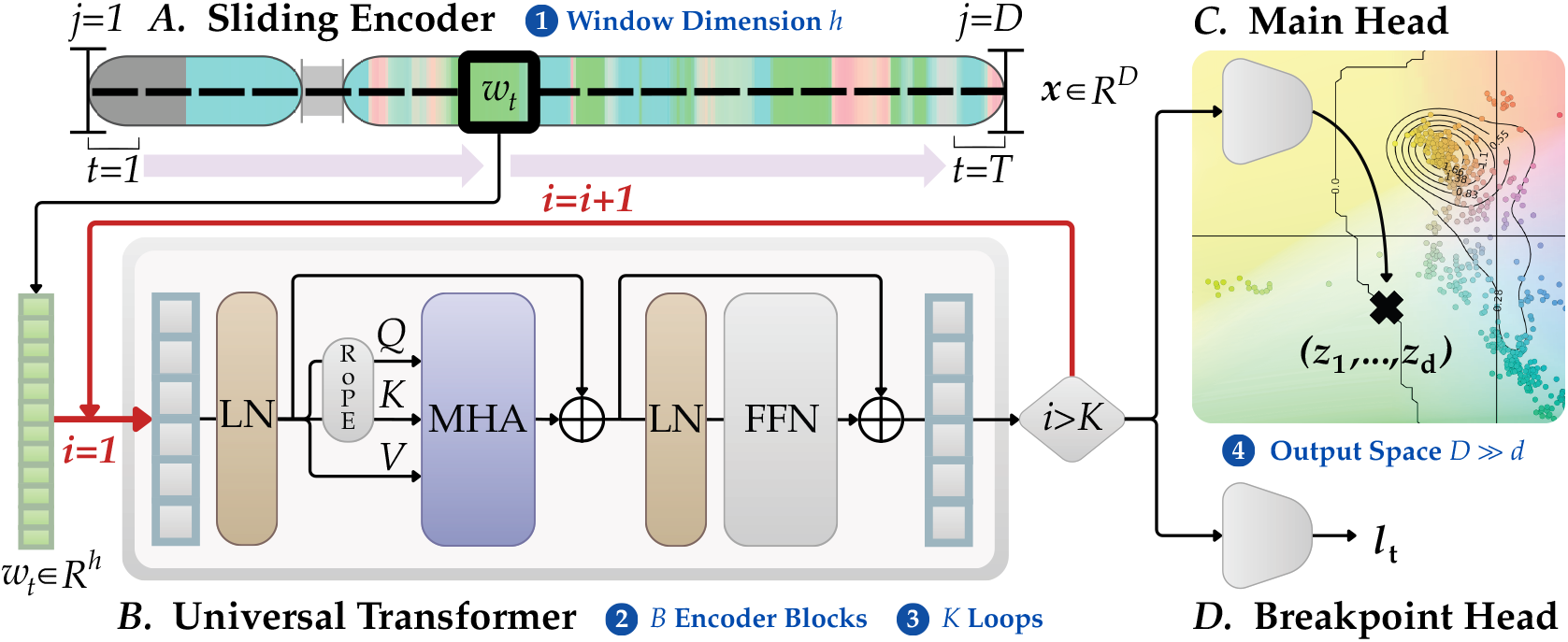
Illustration of the PCLAI model architecture, with four components and three key hyperparameters. Architecture components are listed with letters and hyperparameters are numbered: *A*. Sliding encoder with *(1)* window length of *h* SNPs; *B*. Universal transformer with *B*=2 encoder blocks and *K*=3 self-loops (effective depth is *B* × *K*); *C*. Main coordinate head which predicts the *(4) d*-dimensional coordinates in an output representation space (e.g., PCA), with *D* ≫*d*; *D*. Convolutional breakpoint head that predicts a logit corresponding to the likelihood of being in a window containing a crossover.

On top of these token embeddings we place a transformer encoder with positional structure (**Figure 1.B**). Concretely, we follow the pre-layer norm transformer encoder design [39] for stability at depth and incorporate positional information using rotary positional encoding (RoPE) [40], which applies position-dependent rotations to the query and key vectors and induces a relative position-sensitive dot product in attention. The core of the PCLAI model is a stack of *B* identical transformer blocks. Each block applies layer normalization followed by multi-head self-attention (with RoPE applied inside attention), followed by a residual connection, and then applies a second normalization and a two-layer feedforward network with a Gaussian error linear units [41] and dropout, and applies a final residual connection. We implement a universal transformer-style variant [42] in which, instead of passing through the transformer stack exactly once, we reuse the same stack for a small number *K* of recurrent *self–loops* over the sequence. Effective depth is then the product of the number of layers in the stack and the number of recurrent passes, *B* × *K*. This parameter-sharing across depth can be interpreted as iteratively refining the same representation of each window with a fixed computation block. We chose MLP dropout 0.3, transformer dropout 0.1, a universal transformer architecture with *B*=2 blocks and *K*=3 self-loops (effective depth 6), *d*_model_ = 512, *d*_ff_ = 2048, and 256 as the dimension of the hidden layer in the MLP after grid search hyperparameter tuning (**Supplemental Method S1**).

The resulting per-window representations feed into two prediction heads (**Figures 1.C** and **1.D**). The first, which we refer to as the *main head*, maps each window representation to a *d*-dimensional coordinate vector in the representation space of interest. The second prediction head targets recombination breakpoints. It takes the sequence of per-window representations output by the transformer and applies a temporal convolutional module with multiple kernel sizes and dilated filters. This head integrates information from local neighborhoods of windows at multiple scales and outputs, for each window, a scalar logit that parameterizes the probability that the window contains a breakpoint.

The output is a coordinate-valued mosaic. Because the coordinates live in the reference representation space, one can read off affinities to reference clusters without converting them to hard labels. Many populations are not cleanly captured by a small set of discrete *templates*. A coordinate in a learned space can represent intermediate positions and gradients naturally. Coordinates can be compared across chromosomes and individuals, aggregated, or contrasted with reference haplotypes and assemblies.

### PCLAI objective function

For each phased haplotype, we consider its sequence of non-overlapping SNP windows 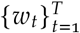 along a chromosome. The model outputs, for each window *t*, a predicted coordinate 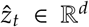 in a chosen representation space and a scalar logit 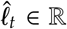 that parameterizes the probability that the window overlaps a recombination breakpoint. In this work, we consider multiple target representation spaces to visualize continuous ancestry. In what follows, we describe the optimization for a genetic embedding space defined by principal components capturing genetic variation of the reference haplotypes, and a geographic representation space defined on the Earth’s surface. In our experiments we also test PCLAI with a genetic embedding space obtained by Uniform Manifold Approximation and Projection (UMAP) [43] with a non-Euclidean distance metric, and the optimization procedure is similar to the PCA case.

In the PCA-based setting, we supervise coordinates against PCA targets 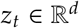 obtained by pooling SNP-level PCA coordinates across the window, and we whiten these targets so that each retained principal component has unit variance under the reference panel. Writing *λ*_1_, …, *λ*_*d*_ > 0 for the eigenvalues of the PCA and 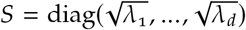, the whitened target is 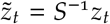 . Formally, let *Z* ∈ ℝ^*d*^ denote the (random) PCA coordinate vector of reference haplotypes in the retained *d*-dimensional PC basis, centered so that 𝔼 [*Z*] = 0. By construction of PCA, the empirical covariance of the reference coordinates in this basis is diagonal: Cov(*Z*) = Λ = diag(*λ*_1_, …, *λ*_*d*_). The whitening transform 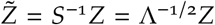 yields an isotropic representation with 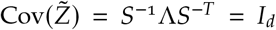 . Whitening prevents the loss terms from being dominated by the highest-variance component and empirically mitigates collapse onto the dominant axis by balancing gradient scales across principal components. Equivalently, whitening converts each PC coordinate into standard deviation units by dividing by 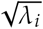, so that a unit error along the *i*-th PC is penalized on the same scale as a unit error along the *j*-th PC, with *i*>*j*. In our experiments we set *d*=3 for PCA-based targets. In the geography-based setting, each window *w*_*t*_ is associated with a direction vector on the unit sphere. The target coordinate *u*_*t*_ ∈ ℝ^3^ is a 3D unit vector (*n*-vector) obtained by mapping a latitude-longitude pair (*ϕ*_*t*_, *λ*_*t*_) to Cartesian coordinates on the unit sphere via *u*_*t*_ = (cos *ϕ*_*t*_ cos *λ*_*t*_, cos *ϕ*_*t*_ sin *λ*_*t*_, sin *ϕ*_*t*_) (with angles in radians) and we enforce the same unit-norm constraint on the model output û_*t*_ by normalizing its three components.

In both cases we denote by *y*_*t*_ ∈ {0, 1} the binary breakpoint label for window *t*, where *y*_*t*_ = 1 if the window overlaps an ancestry boundary and 0 otherwise. To avoid supervising coordinates on windows that overlap breakpoints, we mask these windows in the regression losses. Specifically, we define the valid index set: 𝒱 = {*t* ∈ {1, …, *T*} | *y*_*t*_ = 0} and only include windows in 𝒱 when computing the coordinate loss.

For PCA-based coordinates, the primary coordinate regression loss for a single haplotype is given by a masked *L*_1_ term:

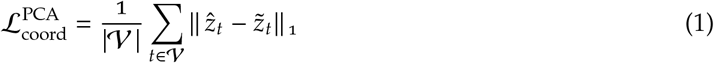

where || · ||_1_ denotes the standard *L*_1_ norm on ℝ^*d*^. This term penalizes deviations in window coordinates along each PCA axis. Because 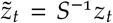, the model is trained to predict coordinates in the whitened space, this is equivalent to a weighted *L*_1_ loss in the original (unwhitened) PC coordinates. In particular, if we map predictions back to the unwhitened PC basis via 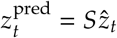, then:

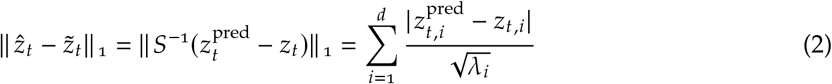

so each PC error is measured in units of that PC’s standard deviation under the reference panel. The geography-based representation is non-Euclidean with the natural notion of distance between two co-ordinates *u*_*t*_ and *û*_*t*_ being the great-circle arc length between their corresponding points on the sphere. For two unit vectors *x, y* ∈ 𝕊^2^ ⊂ ℝ^3^, the angle between them is *θ x, y* = arccos (⟨*x, y*⟩), where ⟨*x, y*⟩ is the inner product; multiplying this angle by the Earth’s radius *R* ≈ 6, 371 km yields the great-circle distance between them. We therefore define the geographic coordinate loss for a single haplotype as:

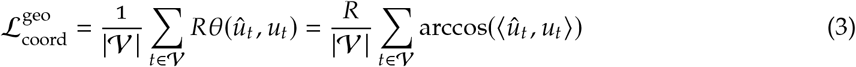

with both *û*_*t*_ and *u*_*t*_ normalized to unit length. This loss is invariant to rotations of the coordinate system on the sphere and has a direct interpretation in kilometers. In practice, we clamp ⟨*û*_*t*_, *u*_*t*_⟩ to [−1, 1] before applying arccos to ensure numerical stability.

To encourage a closer match between the global geometry of the target and predicted point clouds for each haplotype, we add a Chamfer dissimilarity term between the point collections 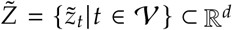 and 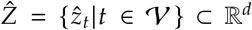 which has proven effective for 3D point-cloud reconstruction in prior work [37]. For PCA-based coordinates, we work in the whitened Euclidean space. This choice admits a precise interpretation in terms of the Mahalanobis distance in the unwhitened PC basis: for any two PC-coordinate vectors *z*_*a*_, *z*_*b*_ ∈ ℝ^*d*^, the rank-*d* Mahalanobis distance with covariance Λ is:

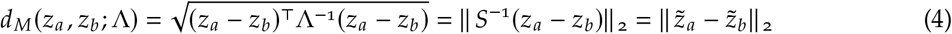

Thus, Euclidean nearest-neighbor matching in whitened space is exactly equivalent to nearest-neighbor matching under a Mahalanobis distance that downweights directions with high reference variance. In this sense, the Chamfer term aligns point clouds using a variance-normalized Mahalanobis geometry induced by the reference panel. The Chamfer dissimilarity between point collections 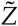 and 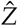 is defined as:

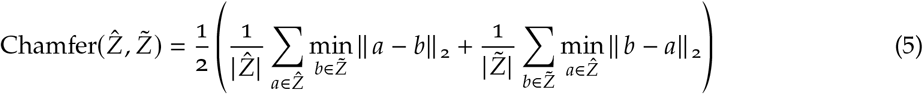

where || · || _2_ is the Euclidean norm on ℝ^*d*^. Intuitively, the first term 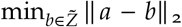 with 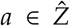 sures that each predicted point lies close to some target point, while the second term, 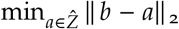 with 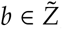, ensures that each target point is approximated by some prediction point. In the geographic setting, we again represent coordinates as unit vectors in ℝ^3^ and compute the Chamfer distance by replacing || *a* − *b*|| _2_ with the great-circle distance *Rθ*(*a, b*) between points on the sphere. We denote the resulting single-haplotype geometric loss by 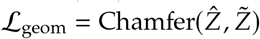 . Unlike the ℒ _coord_ term, the Chamfer dissimilarity is permutation-invariant and therefore acts as a global shape regularizer: it penalizes global discrepancies between the geometries of the target and predicted point clouds without enforcing a one-to-one correspondence between individual windows. This gives us the intuition to weight ℒ _coord_ more than ℒ _geom_ in the joint optimization problem.

Breakpoint supervision is handled via a standard binary cross-entropy loss applied at the window level. Defining 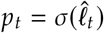 as the predicted breakpoint probability at window *t* with *σ* the logistic sigmoid, the breakpoint loss for a single haplotype is:

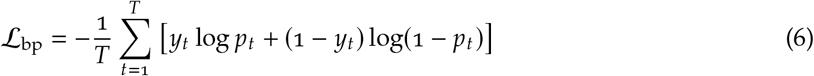

For a minibatch of *M* haplotypes, the overall loss is the weighted sum of the minibatch averages of the three single-haplotype components:

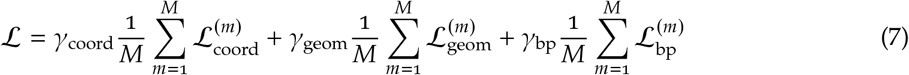

with hyperparameters *γ*_coord_, *γ*_geom_, *γ*_bp_ > 0. In PCA-based experiments we take 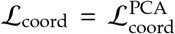, set *γ*_coord_ = 1, and following our intuition we choose *γ*_geom_ < *γ*_coord_ so that the Chamfer term acts as a secondary geometry constraint rather than dominating optimization. In geographic experiments we take 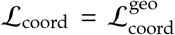, which directly measures the great-circle error in kilometers, and we retain the same structure for *γ*_geom_. We scale *γ*_bp_ so the term contributes a comparable magnitude to the total loss without overwhelming the per-window coordinate regression. In the implementation, the coordinate term is additionally normalized by the coordinate dimension *d*. For optimization, we use adaptive moment estimation (Adam) [44] with learning rate *η* = 10^−3^ and weight decay 10^−5^, and we use early stopping on ℒ. For the specific values of *γ*_geom_ and *γ*_bp_, we used grid search hyperparameter tuning, evaluating the raw loss without scaling each individual loss term. Due to inherent randomness in training, each configuration was run with 5 random seeds. We evaluated all the combinations (**Supplemental Method S1** and **Supplemental Figure S2**); the best-performing weights were *γ*_geom_ = 0.2 and *γ*_bp_ = 10.

### Reference samples and admixture simulation

#### Modern human haplotypes

We assembled two panels of modern human SNP variation. The first panel was designed to capture broad-scale global structure, while the second focused on fine-scale patterns across South Asia (samples from India, Pakistan, and Bangladesh). For the global panel, we used a merge of the 1000 Genomes Project [45] and the Human Genome Diversity Project (HGDP) collections [46]. We restricted to autosomal, biallelic SNPs and removed duplicated markers. We then applied variant filtering with a minor allele frequency threshold of MAF ≥ 0.001. To reduce redundancy while retaining the broad-scale LD structure, we pruned with PLINK 2 [47, 48] using a 50 kb window, a 25 SNP step, and an *r*^2^ threshold of 0.99. Individuals were grouped into subpopulations using the dataset labels. Samples identifying as deriving recent ancestry from a source that could not be assigned to a local geographic region, when in addition that ancestral source was distant from the sampled location, were excluded from training, since no approximate latitude and longitude could be assigned to them for geographic training. These populations included African Caribbean in Barbados, Utah residents with Northern and Western European ancestry and African Ancestry in Southwest US. Additionally, we performed unsupervised genetic clustering [12] at *K* = 7 and retained only those samples with ≥ 95% assignment probability to a single continental ancestry cluster in order to remove from training those individuals with recent transcontinental admixture. Within each subpopulation we split individuals into disjoint training and test sets, holding out 10% of samples as test and using the remaining 90% for training the model and for fitting PCA coordinates. The final panel comprised 2,356 individuals in the training set and 253 individuals in the test set (**Supplemental Table 1**). To construct the target PCA space for PCLAI, we selected the most informative 300,000 autosomal biallelic SNPs from the pruned set by heterozygosity ranking (ranked by 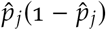, where 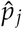 is the allele-frequency estimate for SNP *j* over non-missing genotypes in the reference panel). We retained the first three principal components for supervision and projected both training and test samples onto this space.

The second panel came from the GenomeAsia 100K project [49, 50], which we phased with BEAGLE 5.5 [51] using default settings. We selected individuals with self-reported origins in South Asia, capturing variation across language families, caste and community structure, and recent urban migration (**Supplemental Table 2**). For each subpopulation, we assigned a representative latitude and longitude by selecting the coordinates of a large, densely populated city associated with that population. Because genetic similarity reflects endogamy, historical structure, and migration, we use PCA coordinates as the primary supervision space rather than raw geographic coordinates. To compute these coordinates, we constructed a genome-wide PLINK 2 [48] dataset from the phased panel, restricted to the training individuals, applied standard variant-level QC (MAF ≥ 0.01, missingness ≤ 0.02), performed LD pruning (50 kb windows, 5 SNP step, *r*^2^ ≤ 0.2), and ran PCA to obtain the top principal components. Within this South Asian panel, we curated training and validation sets consisting of 65 subpopulations. We held out a complementary test set of 15 broader or heterogeneous subpopulation labels for test-only evaluation, including umbrella geographic or linguistic labels (e.g., South Indian, Tamil, Gujarati), umbrella social labels (e.g., Scheduled Caste, Sikh), and urban and diaspora samples (e.g., Urban Bangalore). Variants were filtered with the same criteria as the global variation dataset, and the PCA space was built with PLINK 2.

#### Ancient human haplotypes

To explore genetic variation through time, we assembled a panel of ancient and present-day human genomes from the Allen Ancient DNA Resource (AADR) [52], focusing on 1240K capture genomes. We co-analyzed these ancient genomes with modern ones from 1000 Genomes [45] and Human Genome Diversity Project (HGDP) [53] in European, West Asian, and East Asian regions, which had the largest number of samples across radiocarbon dates. We retained individuals with at least 600,000 observed sites and for the analysis, we split the samples into three 1,000-year historical bins and one modern era bin, specifically: the Late Bronze/Iron Age 1500 BCE–500 BCE, Classical Antiquity period 500 BCE–500 CE, the Medieval Period 500 CE–1500 CE, and finally the Modern Era 1500 CE–2000 CE (**Supplemental Table 3** and **Supplemental Figure S1**). Variants were restricted to autosomal, biallelic SNPs. Because the ancient callset was aligned to the GRCh37 reference genome whereas our 1000 Genomes and HGDP reference panel was on GRCh38, we lifted over the ancient VCF and sorted and normalized alleles against the GRCh38 reference genome using CrossMap [54] and BCFtools [55]. We performed phasing and imputation using BEAGLE 5.5 [51, 56]. Prior work [57, 58] suggests that broader reference panels can improve imputation accuracy compared with restricting the reference to a single continental subset, so we impute using the full modern reference panel described above.

For the geographic target, we associated each subpopulation with the approximate latitude and longitude at which the ancient DNA sample was found. We operate under the assumption that in ancient times long range lifetime migration was less frequent (**Supplemental Results S1**). For the PCA representation, we first intersected the AADR subset sites with our modern human genome sites (**Supplemental Table 1**) which resulted in 793,962 biallelic SNPs. We then applied QC and LD pruning with PLINK 2 [48] (MAF ≥ 0.05, 200 kb windows, 50 SNP step, *r*^2^ ≤ 0.2) and selected 260,000 most informative SNPs according to heterozygosity ranking. As an additional QC signal for downstream analyses, we recorded the pre-imputation missingness, and we constructed a PCA space with modern references (selecting the most informative 260,000 autosomal biallelic SNPs) and projected the ancient samples that had <25% missingness (**Supplemental Figure S6**).

#### Admixture simulation from reference haplotypes

To obtain ground truth breakpoints and ancestry coordinates at realistic recombination scales, we simulated admixed haplotypes by reassembling mosaics of the phased reference haplotypes. Using a forward-in-time simulation, we take a batch of real haplotypes, attach continuous descriptors (e.g., PCA or geographic coordinates), and then recombine the segments across haplotypes according to the Wright-Fisher recombination model [59], with a number of generations sampled uniformly at random for each batch, which loosely sets the expected number of crossover events. Each shuffle creates a new mosaic haplotype with a full record of where ancestry switched and what the local coordinates should be in each window. For each chromosome, we sample the number of crossovers as *K* ∼ Poisson (*λ*) with *λ* equal to the chromosomal genetic length in morgans, then sample breakpoint locations according to the sex-averaged recombination map from [60]. We precompute per-SNP recombination rate by interpolating centimorgan positions and take a finite difference. We choose the starting parental haplotype with probability ½ and generate the transmitted haplotype by alternating between the two parental haplotypes at successive breakpoints, recording the resulting segment boundaries as ground-truth crossover locations.

For PCLAI, we need window-level targets rather than per-SNP descriptors. We therefore partition the SNP axis into non-overlapping windows of fixed size *w*_*t*_ = 500 or 1,000 variants and pool labels within each window with an approximate mode for continuous values. The function computes, for each window of SNPs, a small histogram over a fixed number of bins (we use 32 by default), finds the most populated bin, and returns its midpoint as the approximate mode. Applying this function yields a window-level coordinate that reflects the densest cluster of SNP-level coordinates, rather than a straight average. In practice, this reduces the “blurring” of coordinates. This mode pooling is especially important at realistic crossover scales, where a small number of breakpoints can fall inside a fixed SNP-count window and would otherwise pull the target coordinate toward an artificial intermediate.

Breakpoint labels are constructed in parallel. At the SNP level, we mark a breakpoint wherever the discrete label changes between consecutive positions along a haplotype. This produces a boolean matrix indicating sites of change at single-SNP resolution. We then aggregate this mask over windows: a window is labelled as containing a breakpoint if *any* SNP in that window shows a change. This produces supervision that matches the model’s operational goal, i.e., detecting whether a window overlaps a recombination junction, while leaving precise sub-window localization to the continuous coordinate signal. For our experiments, we simulate 160,000 admixed haplotypes for training and 48,000 for testing, each using up to 64 generations of simulated recombination. Varying the number of generations modulates the breakpoint density and expected segment length, providing training examples that span a realistic range of ancestry tract lengths under recombination [59]. For modern references we use 1,000 SNP windows and for ancient references we use 500 SNP windows due to differences in the number of sites available.

## Results

### A geometric surrogate for genetic distance and admixture

Mapping genotypes directly onto geographic latitude and longitude can be informative in some isolation-by-distance settings, such as modern Europe [19], but it does not generalize across the world or time, due to migration and endogamy (**Figures 2** and **3**). A clear example is seen in the Indian subcontinent (**Figures 4** and **5**), where genetically distinct groups often live in the same exact location, due to endogamy breaking the local panmixia assumption which underlies gene-geography correspondence. India is not unique in this respect; in Europe smooth gene-geography gradients across the continent are in part a construct of investigator-driven population selection schemes. Genes would not mirror geography so strikingly if all Europeans, for instance Roma, Ashkenazi Jewish and Sami groups, were included. Genetic ancestry also changes geographically over time, making geography a confusing label for genetic ancestry [2].

**Figure 2.**
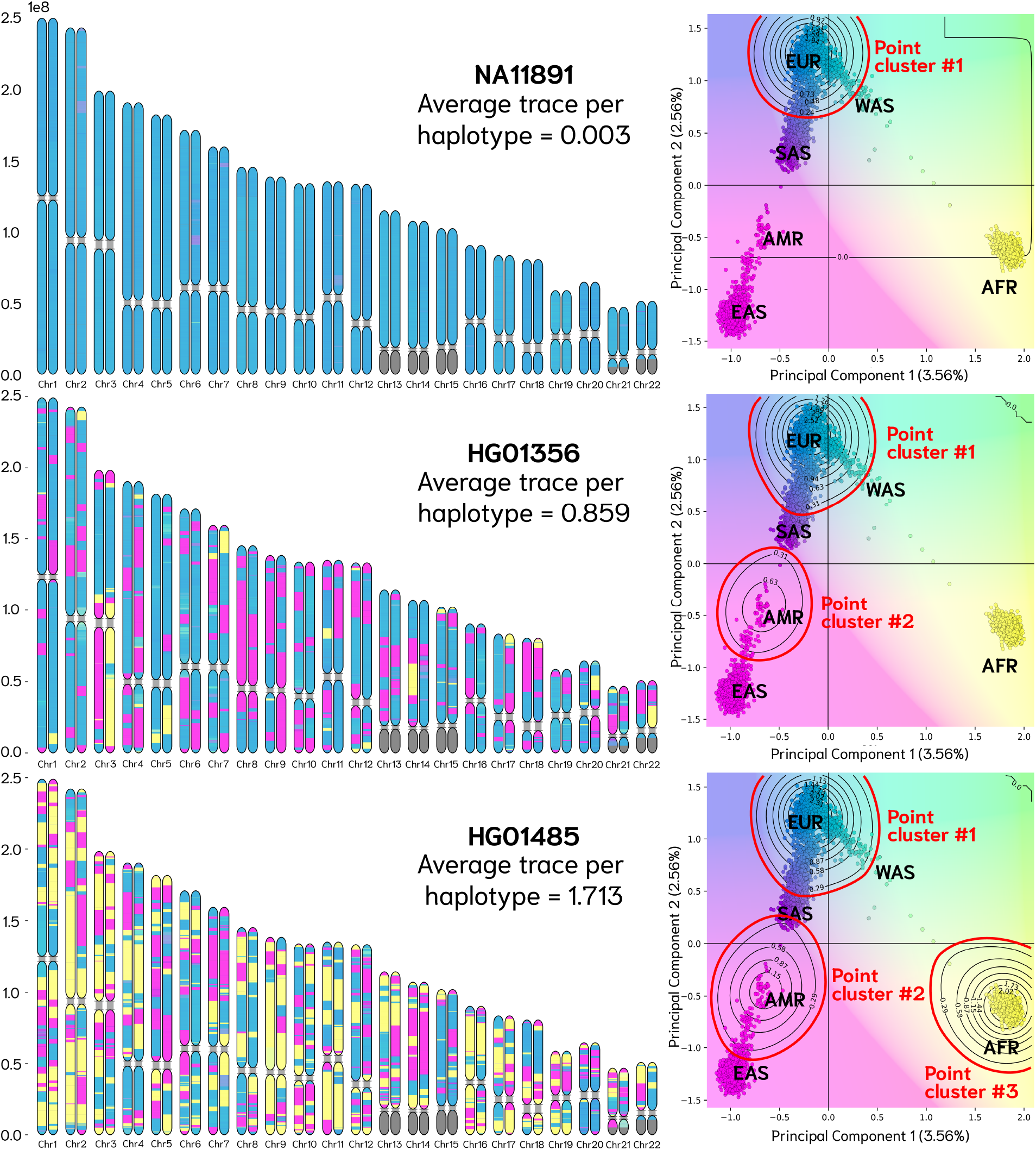
Comparison of samples with minimal, intermediate, and maximal average within-haplotype point-clouds Tr (Σ). *Left* panels show haplotype-level ancestry paintings for the 22 autosomes, illustrating increasing within-haplotype admixture from top to bottom. *Right* panels show the corresponding 2D PCA projections, where references are shown as colored points and the density of low-breakpoint probability predictions is overlaid as a contour map. The average point-cloud Tr (Σ) (total variance) increases monotonically across panels, reflecting progressively more dispersed and elongated point clouds, consistent with increasing structural and ancestry complexity: (*Top*–*Bottom*) sample NA11891 (Utah residents with Northern and Western European ancestry), sample HG01356 (Colombian from Colombia), and sample HG01485 (Colombian from Colombia).

**Figure 3.**
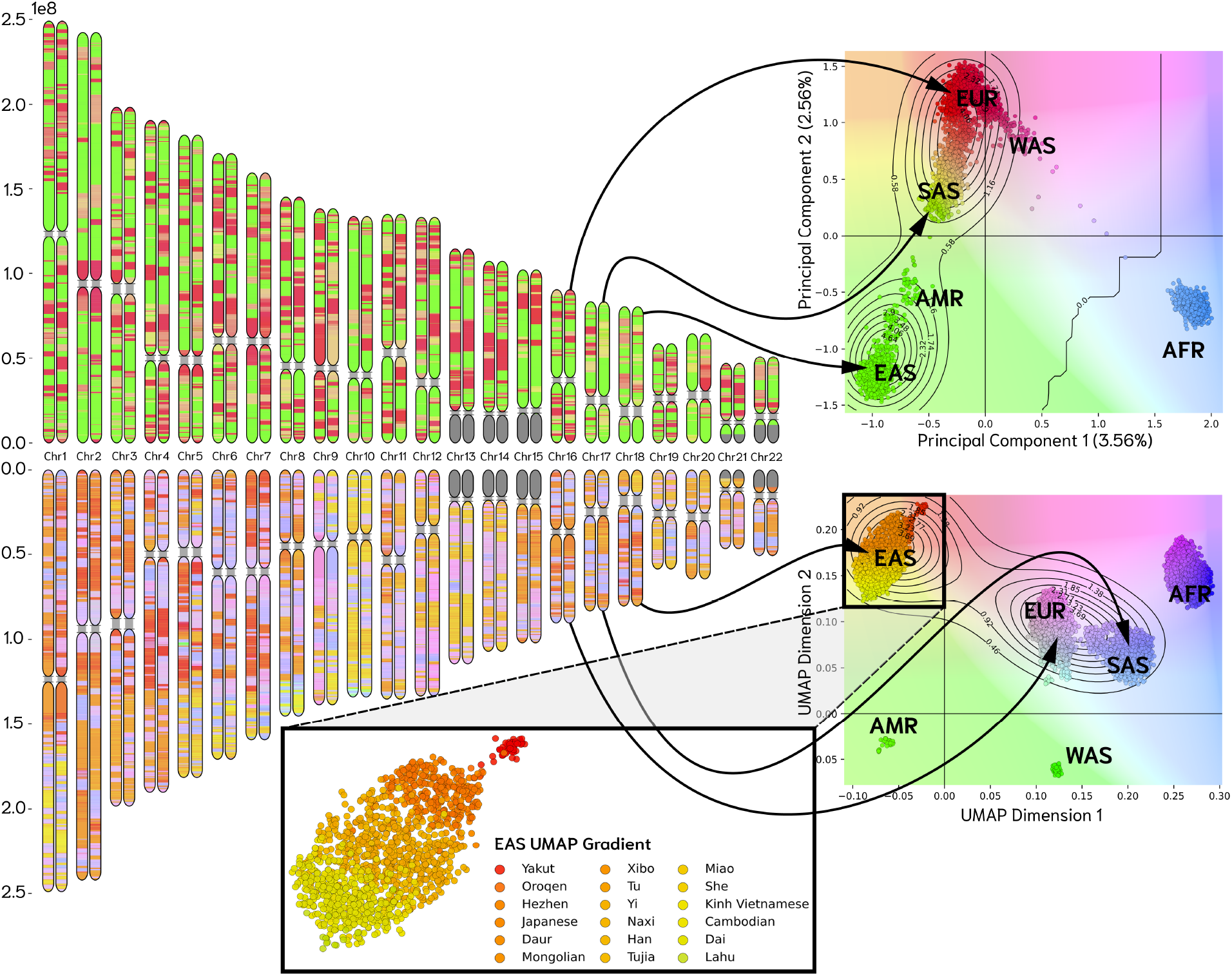
PCLAI chromosome paintings for HGDP00099 (Hazara in Pakistan) highlight robust break-point detection under PCA and UMAP embedding spaces. *Left*: haplotypic PCLAI paintings (Chromosomes 1–22) for HGDP00099 (Hazara in Pakistan), showing window-level ancestry coordinates along each chromosome. *Right*: the corresponding predicted point clouds overlaid on the reference embedding, shown in (*top*) PCA space and (*bottom*) non-Euclidean UMAP space with reference individuals plotted as colored points and low-breakpoint-probability predictions summarized by density contours. Curved arrows link representative chromosomal segments to their locations in the embedding spaces, illustrating that while the geometry of the coordinate space changes between PCA and UMAP, the inferred ancestry mosaics and junction patterns remain visually consistent across embeddings.

**Figure 4.**
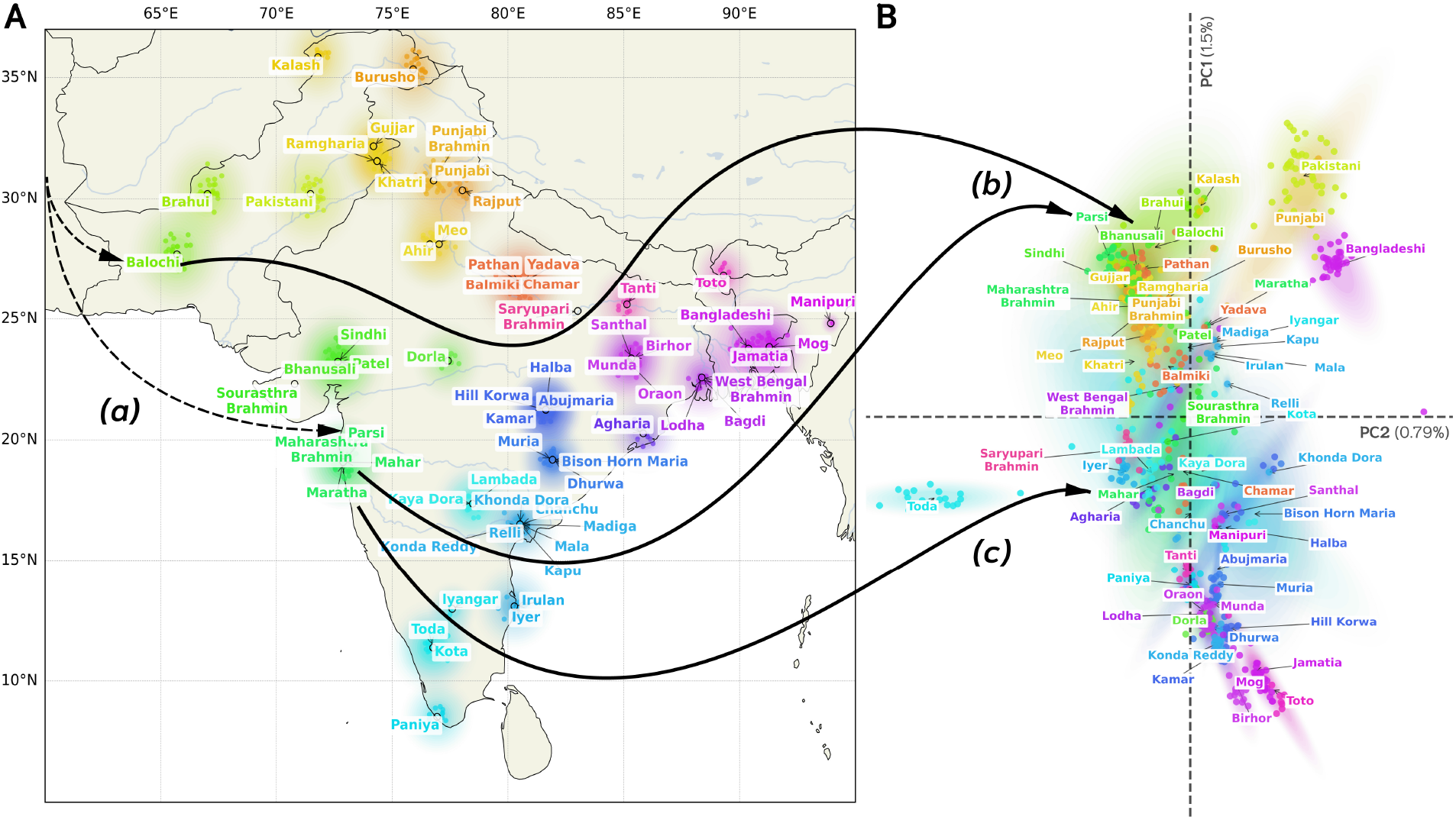
Genes do not always mirror geography. *Left*: geographic map of South Asian populations from GenomeAsia 100K [49, 50]; *Right*: the same populations in genetic PCA space with percent variance explained by each axis indicated. *(a)* highlights three example subpopulations: Balochi, Parsi, and Mahar. Balochi and Parsi are separated geographically but share substantial Iranian-related roots, whereas Mahar is geographically close to Parsi, but shifted toward more ASI-enriched groups in the genetic space. In the PCA, *(b)* Balochi and Parsi cluster near one another and near other Iranian-related groups such as Brahui, while *(c)* Mahar falls along the Indian cline toward central/southern groups, illustrating how geography and genetic structure decouple under the combined effects of stratified ancient admixture and subsequent endogamy.

**Figure 5.**
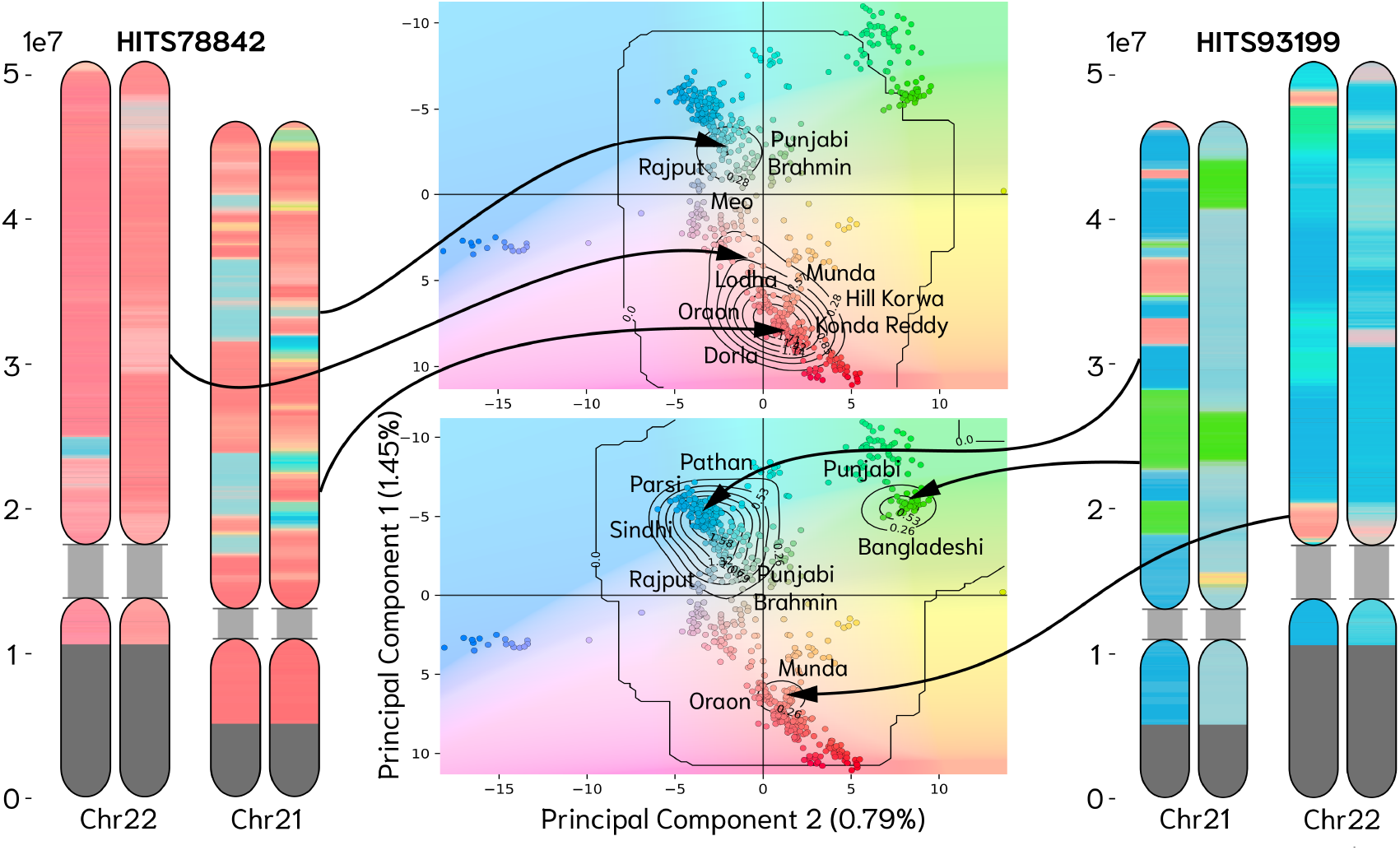
PCLAI on South Asian genomes uncovers ANI-ASI variation at a local level. The *central* panels display the reference genomes in the PCA space together with kernel-density contours summarizing the distribution of genomic window-level predictions for each individual; the straddling chromosome capsules correspond to haplotype-resolved paintings along Chromosomes 21 and 22. Colors along the chromosomes encode the predicted local PCA coordinate for each window. Abrupt color transitions mark recombination breakpoints between ancestry segments that occupy measurably different positions in the PCA space.

For these reasons, we do not initially use geography-based coordinates and instead use principal components of the whole genome samples to construct a geometric space for annotating local ancestry. Unlike latitude and longitude, distances in this coordinate space have a natural genetic interpretation. In particular, because the *F*_2_ statistic of Patterson et al. [2], which is a measure of accumulated genetic drift or branch length between populations, is the expected squared Euclidean distance between their allele-frequency vectors, and because the principal components define a lower dimensional space that minimizes the squared residuals of projected points, distances between points in this PCA space can be interpreted as approximations of their genetic distance (*F*_2_) [61]. This geometric interpretation is further supported by a genealogical perspective [5], as SNP projections onto principal components can be obtained by average coalescent time for pairs of samples.

Consider a phased haplotype, whose each genomic position is labeled with a coordinate *x* ∈ ℝ^*d*^ in the PCA space; the set 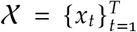 defines the point cloud representing its local ancestries. The mean of the point cloud is defined 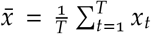 and covariance 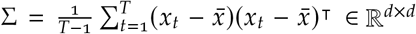 . Because the squared Euclidean distance in PCA space approximates *F*_2_, the trace of the covariance matrix 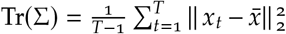 is an average of the within-haplotype *F*_2_s across the genomes. (This is because 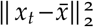 computes the distance between the local ancestry assignment, and the mean haplotype PCA coordinate.) In other words, when different genomic segments trace back to sources separated by substantial drift, the point cloud must also spread out in the vector space. Thus, Tr Σ gives a measure of the point-cloud spread that can be used to quantify the extent of admixture spread across genetic ancestries (**Figure 2**). This point cloud view also clarifies why local ancestry remains an informative approach while, in contrast, a global PCA placement can lose visible admixture signal after relatively few generations, even though ancestry tracts along the genome remain well-defined mosaics [5]. If we want to elaborate further on point cloud topology, we can examine the eigenvalues of Σ. Let *λ*_1_ ≥ *λ*_2_ ≥ … ≥ *λ*_*d*_ > 0 be the eigenvalues of the covariance matrix, then if Tr (Σ) is small and the eigenvalues are close to zero, we can conclude that the point cloud is tight and the local ancestries are very similar. However, if Tr (Σ) is moderate and we have *λ*_1_ ≫ *λ*_2_, …, *λ*_*d*_ this will result in an elongated scatter along a single axis suggesting that ancestry varies mainly along one drift direction. If we observe multiple *λ*_*i*_ that are large, then this signals that the admixture is multi-directional, and we have an admixture of haplotypes from multiple independently drifted ancestries.

### Chromosome paintings across coordinate spaces

PCLAI is versatile, adaptable to any coordinate space as output for local ancestry regression. If break-points are real biological junctions, they should be stable across different embedding spaces, and we test that invariance: in **Figure 3** we show the predictions for the same sample HGDP00099 (Hazara in Pakistan) using two different spaces as outputs. In the first case, we predict for each genomic position a coordinate in the PCA space; in the second case, we predict coordinates in the UMAP space [43]. Even though the coloring is different, because the coordinates live in different embeddings, the locations of color changes—and thus the inferred recombination breakpoints—are visually consistent. Within the UMAP-based painting, we observe an East Asian (EAS) gradient (yellow-green ↔ red) for segments that fall inside the EAS-like manifold. UMAP distorts global geometry, but highlights subtle variation less apparent in the PCA embedding with windows colored toward yellow-green localizing nearer South-east Asian references (e.g., Dai in China, Cambodian in Cambodia, Lahu in China), orange tones falling closer to more northern East Asian references (including Han Chinese in China and neighboring EAS groups), and red tones that are closest to Northeast Asian/Siberian-indigenous-like references (e.g., Yakut in Russia, Oroqen in China). In contrast to discrete LAI, PCLAI allows capturing fine-grained details such clinal gradients; for clarity, the “EAS UMAP gradient” zoom-in is discretized relative to the UMAP panel, assigning discrete distinct colors along the EAS-like gradient. Also note that the PCA space is governed by the Euclidean distance, and the UMAP we used is fitted using the Hamming metric (and 10 neighbors, minimum distance 0.5); essentially, we use different embedding spaces and metrics, but there is consistency in breakpoint detection. An additional example of an African Ancestry in South-west US sample is provided in **Supplemental Result S2** and **Supplemental Figure S4**.

To move beyond this single chromosome example and quantify agreement between coordinate spaces, we compared window-level breakpoint probabilities for the PCA-based and UMAP-based PCLAI models across all 654 test genomes (**Supplemental Table 1**). For statistical testing we treated the genome (41, 972 windows) as the unit of inference and summarized agreement per sample. Across samples, Pearson correlation between breakpoint probabilities in the two models was moderate to high ( 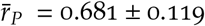 one-sample *t*-test for 𝔼 [*r*_*P*_] >0: *t* (653) ≫ 1, *p*<10^−15^), while Spearman correlation was even higher 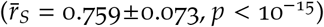 . The higher rank correlation indicates that the two models are especially consistent in the *ordering* of windows from least to most breakpoint-like, even if their probability scales differ slightly. We then thresholded breakpoint probabilities at *p*≥0.5 and compared binary break-point calls. Chance-corrected agreement, measured by Cohen’s *κ*, was substantial 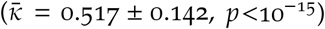, and the mean Jaccard overlap between the sets of breakpoint windows was 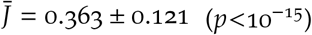 . Given that true recombination events are sparse along the genome, a Jaccard index of this magnitude implies that a large fraction of windows flagged as breakpoints by either model are shared. Finally, the mean signed difference in breakpoint probability between the PCA and UMAP models was very small ( 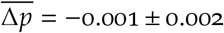 two-sided test of 𝔼 [Δ*p*] =0: *p* = 2 ×10^−35^), indicating negligible systematic bias. Taken together, these results suggest that PCLAI produces robust and largely interchangeable breakpoint predictions across very different embedding space objectives—one Euclidean distance, and one based on the Hamming metric. PCA and UMAP emphasize different details of the coordinate geometry, but they highlight essentially the same genomic regions as ancestry junctions.

### Navigating South Asian genetic structure with PCLAI

Ancient DNA and genome-wide studies now agree that most present-day South Asians descend from an admixture of at least three different sources: Indigenous South Asian hunter-gatherers (AASI), Iranian farmers from the Indus Periphery cline, and Steppe-derived pastoralists that also contributed to Bronze Age Eastern Europeans [62, 63]. Across diverse Indian and neighboring populations, the first principal component typically recapitulates the cline that emerged from the admixture between ancestries related to ancient West Eurasian–associated sources (Ancestral North Indian, or ANI), which themselves largely reflect mixtures of Indus Periphery–related groups with Steppe pastoralists; and deeply rooted South Asian ancestries (Ancestral South Indian, or ASI), which are modeled as mixtures of Indus Periphery– related groups with additional AASI-enriched ancestry. The differential admixture of these groups across Indian subpopulations was then followed by long periods of within group endogamy that amplified intra-community drift and propagated between-group differences [62, 63, 64, 65]. In **Figure 4** we compare the geography of the Indian subcontinent and the PCA of GenomeAsia 100K genomes [49, 50] sampled from that region. We can clearly see that the mapping from geography to genetic space is not simple. In the PCA, populations from the geographic north and northwest (e.g., Pathan in Pakistan, Sindhi in Pakistan, Punjabi Brahmin, Rajput) occupy the high-ANI and Steppe-enriched end of the first principal component, while many central and east Indian Adivasi (or tribal groups) and Austroasiatic-associated groups (e.g., Munda in Bihar, Oraon in Bihar, Hill Korwa in Jharkhand) shift toward the opposite end of the axis, consistent with increased ASI- and AASI-related ancestry [62, 64, 65].

Within this coordinate system, **Figure 5** shows chromosome paintings for two GenomeAsia 100K v2 individuals. The visualization makes explicit a key benefit of coordinate-space regression: both the dominant ancestry mode (where most windows fall) and secondary modes (small but consistent excursions toward other regions of the PCA) can be read directly from the density contours and then localized to specific genomic tracts on the painted chromosomes. The Gaud in Odisha individual (HITS78842)— sampled in the Sundargarh district—provides a clear example of a genome-wide projection near central and east Indian Adivasi and tribal-belt populations, where the dominant mode is mirrored on both Chromosomes 21 and 22 by long stretches of relatively similar coloring mapping to groups from the Munda–Oraon corridor and adjacent populations (e.g., Tanti in West Bengal, Oraon in Bihar, Munda in Bihar, Muria in Chattisgarh, Hill Korwa in Jharkhand, Lodha in West Bengal, Dhurwa in Jharkhand, and Kamar in Chattisgarh), and showing additional affinity to other south and central Indian ASI-enriched groups such as Paniya in Kerala and Konda Reddy in Andhra Pradesh. This pattern is consistent with previous work [64, 65] that modeled many Adivasi and tribal groups as having high proportions of ASI and AASI-related ancestry with minimal Steppe contribution. In contrast, the sample lies far from high-ANI reference clusters and exhibits short, scattered tracts whose predicted coordinates move toward the northwest Indo-Aryan references (e.g., Rajput, Meo in Haryana, and Punjabi Brahmin). In admixture terms, this pattern is consistent with a predominantly ASI-enriched ancestry profile with a small northwestern component that is visible as a weak but detectable pull along the first principal component. Importantly, in Indian population history such minor pulls can reflect broadly shared West Eurasian–related ancestry that is widespread at low levels across many South Asian groups.

By contrast, the Sikh individual (HITS93199) from Punjab in **Figure 5** has local coordinates concentrated in the northwest South Asian portion of the genetic space with only limited excursions toward other regions. The window-level contour is tightly centered on the Punjab–Sindh–northwest India and Pakistan corridor, overlapping reference clusters such as Punjabi Brahmin, Rajput, Sindhi in Pakistan, and Pathan in Pakistan, and extending toward other West Eurasian-rooted South Asian references (e.g., Parsi). This is consistent with previous findings that many northwestern groups carry elevated proportions of Steppe- and Iranian-farmer–related ancestry compared with more ASI-enriched groups farther along the cline [62, 65]. A smaller set of rare windows approach the central and east Indian Adivasi-associated region (Oraon/Munda corridor), reflecting the pervasive but heterogeneous imprint of ANI– ASI mixture that has been detected in almost every Indian group surveyed [62]. Taken together, these two case studies demonstrate how PCLAI leverages the geometry of South Asian PCA to produce interpretable, haplotype-resolved mosaics that align with established demographic models of Indian genetic variation while adding locus-specific resolution. In **Supplemental Figure S5** we present chromosome paintings of 4 Nanabuddha samples from Maharashtra and 4 Scheduled Caste samples from Gujarat, both labelled as Dalit groups in the varna system [66, 67]. Even though they are labeled under the same umbrella group, we show that both groups present very differentiated genetic ancestry paintings.

### Exploring ancestry through time with PCLAI trained on ancient human genomes

To probe how admixture in the past shaped present-day human genomes, we trained four time-stratified PCLAI models using ancient and modern genomes binned by historical period. Specifically, we use human AADR genomes [52] extracted from skeletal material for the Late Bronze and Iron Age (1500–500 BCE), Classical Antiquity (500 BCE–500 CE), and the Medieval Period (500–1500 CE), and we additionally included 1000 Genomes [45] and HGDP [53] genomes to define the Modern Era (1500–2000 CE) panel.

Applying an ancient-trained PCLAI model to a modern individual results in a chromosome painting that shows the most genetically similar ancient mosaic of haplotypes to the modern sample. In other words, each modern haplotypic segment is mapped to the coordinate where the most similar ancient haplotypes from that time bin concentrate. To make the insights on ancestral population movements as direct as possible, we trained these time-stratified models on geographic coordinates rather than PCA and used them to “time-travel” paint a modern genome. Below we illustrate our time exploration of the 1000 Genomes sample HG00140 (British in England and Scotland) with the results summarized in **Figure 6**.

**Figure 6.**
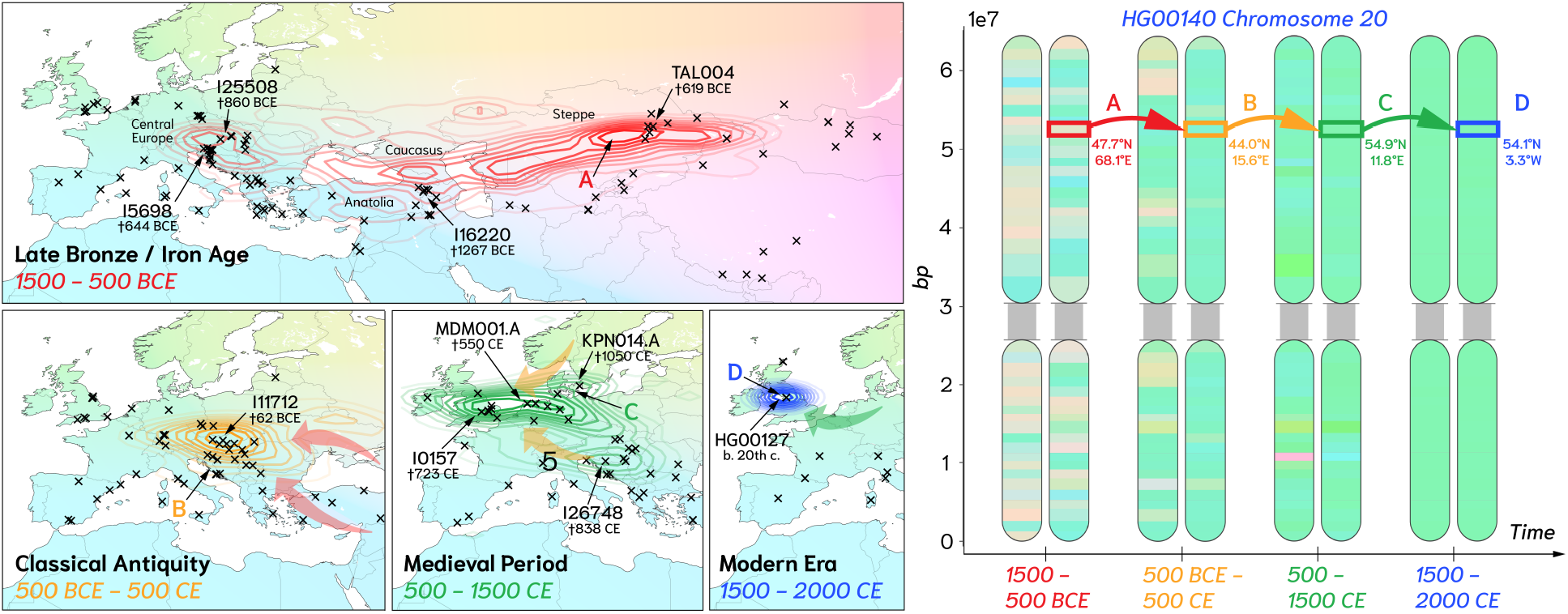
PCLAI trained on time-stratified ancient genomes allows for “time-travel” chromosome paintings. *Left*: for each historical training bin (Late Bronze/Iron Age, Classical Antiquity, Medieval Period, Modern Era), kernel density contours summarize the geographic coordinates inferred for HG00140’s point cloud, overlaid with the locations of training individuals (black crosses) and annotated representative ancient samples located and discussed in the text. *Right*: Chromosome 20 paintings for HG00140 under each time-stratified model (two haplotypes per time bin), illustrating how local inferred coordinates vary along the chromosome. Colored rectangles track the same genomic interval across time bins, highlighting a trajectory in inferred coordinates (A–D; latitude/longitude shown) from steppe-associated locations in the oldest model toward northwestern Europe in the Migration Period and ultimately the United Kingdom in the Modern Era.

We start in the Modern Era model (1500-2000 CE). Here, the point cloud mode for the painted genome falls in the United Kingdom (**Figure 6**), which aligns clearly with the documented geographic origin of this British sample. Stepping back to the Medieval Period model (500–1500 CE), the dominant point cloud mode shifts towards the Anglo-Saxon-Jutland/North Sea region—precisely the zone that was inhabited by Germanic tribes who migrated from what is now Denmark, the Netherlands, and northern Germany into Britain. The map panels also display (as crosses) the ancient individuals used for training, and representative medieval samples from northwestern Europe [68] appear near the modes of the inferred point clouds. One such sample is the medieval tooth KPN014.A (†1050 CE) found on Danish soil; [68] report that it shares most alleles with modern Swedish and Baltic populations, indicating an influx from a Scandinavian source ancestry as well. Another representative sample is the MDM001.A (†550 CE), a genome recovered from a tooth in the Netherlands, from Friesland—an area from which many inhabitants embarked for Sub-Roman Britain as part of the Anglo-Saxon migrations to England following the 476 CE collapse of the Western Roman Empire. A further anchor point for HG00140 in this period is a tooth I0157 (†723 CE) from South Cambridgeshire, recovered from a site that came to light when pasture land was first ploughed and cultivated, revealing early Anglo-Saxon burials [68]. Alongside this primary northwestern European mode, we also observe a smaller point cloud mode located in Croatia, with an affinity to the genetic material from the petrous skeletal element I26748 (†838 CE) from an Early Medieval necropolis. [69] describe this individual’s ancestry as an admixture between Anatolia Roman and Eastern Europe; analogous to the Germanic influx in post-Roman Britain, the early medieval Balkans saw substantial post-Roman migration.

Classical Antiquity (500 BCE–500 CE) reveals a different picture. Under the model trained on ancient genomes from Classical Antiquity, the largest point cloud mode falls in the Middle Danube area, which for much of this interval was governed by the Roman Empire. [69] emphasize that the Danubian frontier was a zone of constant military activity and exchange with populations living north of the frontier including Celts and Goths—an interpretation that helps connect the Classical Antiquity signal to the stark northward shift we see once we move into the post-Roman Medieval model. It is reported that various Germanic (Gothic) groups settled in the Danubian region alongside other non-Roman groups such as Celts and later the Huns and Slavs. A representative sample for this Classical Antiquity mode is I11712 (†62 BCE), extracted from petrous bones found below Roman and Celtic pottery fragments and near Celtic-Roman architectural remains [70].

The Late Bronze/Iron Age model (1500–500 BCE) reaches further back into a period when [71] report a major ancestry shift occurring (<900 BCE); this concurs with the strong point cloud warp we observe in this oldest time-stratified painting (**Figure 6**). Historically, this period coincides with major technological developments that increased mobility: the prior development and adoption of draft-powered transport in the Near East and Pontic-Caspian Steppe had enabled movement, including along the Southern Arc (bridging Europe through Anatolia with West Asia) [71, 72]. In our model’s point cloud, this shows up as a prominent Steppe-related ancestry signal—explicitly including Pontic-Caspian and western Steppe influences from the broader Kazakhstan region. Western and central European affinities remain detectable in the point cloud: we capture Celtic influences through I25508 (†860 BCE), a cranial fragment found in a burial settlement in Hungary, but the overall density mass shifts primarily toward the Caucasus and Eurasian Steppe. Another close representative sample is I5698 (†644 BCE), a petrous bone from an Early Iron Age cemetery in Slovenia associated with the Hallstadt culture, one of the founders of the Celtic tradition. A representative Late Bronze Near East sample found near the mode is I16220 (†1267 BCE), recovered from an Armenian Late Bronze Age cemetery. [72] report analyses of Late Bronze Age samples from Armenia, showing ancestry flowed from the steppe not only west of the Black Sea into Southeastern Europe but also across the Caucasus into Armenia. Finally, one of the most representative individuals for the Late Bronze/Iron Age point cloud is TAL004 (†619 BCE), found in a burial site in Central Kazakhstan; its funerary structures were similar to those of the Iron Age Tasmola culture [73], which arose from the nomadic Eastern Scythians.

This framing also clarifies a natural question: why do we not see a “Britannic” match for HG00140 under the Iron Age model? The simplest interpretation is that the people inhabiting Britain in that period were genetically quite different from this modern British sample, so the closest geographic-genetic matches for HG00140’s segments (when constrained to Late Bronze/Iron Age training data) dwelt elsewhere in Eurasia at that time. Empirically, this aligns with existing evidence that there was comparatively less gene flow from continental Europe during the Iron Age and a relatively independent genetic trajectory in Britain [70]. Later, migrations into Britain likely contributed most of the haplotypes found in this individual. Indeed, even in the historical period we know the Romano-British population was partially replaced by migrants from the Germanic-speaking part of the continent [68]. In this sense, the absence of a tight Britain-centered mode in the oldest time slice is exactly the kind of temporal geographic difference that time-stratified, coordinate-based local ancestry painting is meant to surface. And this pattern that present-day inhabitants of a region are often a poor proxy for the populations that lived there in the distant past has been observed in multiple other instances (**Supplemental Result S3**) [3, 74, 75].

Another observation is that as we condition on progressively older time bins, the inferred chromosome paintings become more and more admixed and the corresponding point clouds become broader (**Figure 6**). One contributor is purely genealogical: the number of ancestors in the pedigree grow roughly exponentially with time, while the expected length and detectability of shared haplotypic segments inherited from any particular ancestor decay with recombination [4]. As a result, beyond a modest number of generations, any single descendant or kin genome is expected to retain only a sparse and uneven subset of its distant ancestors’ genomes, and segment-level matches to a finite reference panel become increasingly sensitive to stochastic variation in recombination. In the geographic PCLAI setting, this manifests as a diffuse set of coordinate affinities for older periods, a broader ancestral point cloud and more locally variable paintings, emphasizing the point that all human are admixed when considered from the standpoint of reference haplotypes at some time in the past.

## Discussion

PCLAI reframes local ancestry inference from a classification problem to a point cloud regression problem. Instead of labeling each genomic segment with one of a small, incomplete set of arbitarily discretized ancestry categories (that in many cases are partially socially constructed and represent an imperfect proxy for genetic similarity [6, 23]), we infer a continuous coordinate for each haplotypic window in an embedding space than can be chosen to be based on genetic similarity. Separately, we estimate the probability that the window overlaps a recombination breakpoint. This results in an ancestry representation that is both continuous within haplotypic segments and explicitly piecewise through recombination—an output that is naturally expressed as a point cloud indexed along the genome. By operating directly in continuous spaces, PCLAI avoids forcing clinal and intermediate ancestries into imperfect categorical templates, while preserving the ability to localize ancestry variation to specific chromosomal segments.

The point cloud framing supports quantitative summaries of admixture geometry in addition to visualization. In PCA space, squared Euclidean distances approximate drift distances and average coalescence time [2, 5], motivating within-haplotype dispersion statistics such as Tr (Σ) as interpretable measures of how widely a haplotype’s local ancestries spread in the embedding space. In practice, we observe that genomes with minimal recent admixture with respect to the references used to span the embedding space produce compact, unimodal point clouds, whereas more complex admixed genomes produce dispersed and often multi-modal point clouds with localized segments corresponding to excursions toward distinct regions of the reference embedding. This perspective distinguishes global ancestry mixtures from locus-specific chromosome paintings, providing a unified language for describing both.

Comparing an Euclidean PCA objective to a UMAP objective fitted with a Hamming metric, we find that breakpoint probability landscapes remain substantially concordant across models. At the same time, the coordinates themselves retain embedding-specific meaning: PCA coordinates emphasize major drift axes, whereas non-linear embeddings can emphasize local neighborhoods or alternative similarity metrics. The separation between stable breakpoints alongside embedding-dependent coordinates suggests a practical strategy for analysis: treat breakpoints as a relatively invariant discrete signal and choose the embedding space based on the scientific question and interpretability needs.

The South Asian example provides a canonical instance where discrete ancestry assignments can be limiting and cannot capture the full richness of language-, geography-, and community-structured demographic history. In this context, PCLAI leverages the geometry of the South Asian PCA space to produce haplotypic mosaics that make both dominant ancestry modes and secondary components visually and quantitatively accessible. This capability is particularly valuable when ancestry variation is distributed along a relatively small number of dominant axes yet remains heterogeneous across the genome due to historical mixture, subsequent drift, and community structure. Training PCLAI on time-stratified ancient panels further highlights that ancestry is not a fixed attribute tied to present-day geography: the same modern genome can receive systematically different coordinate paintings depending on the time period used to define the reference representation space. In our ancestry-through-time analyses, geographic-coordinate models trained on different historical bins produce point clouds whose modes shift across the continent in patterns consistent with documented ancestral population movements, while also revealing discontinuities—such as the absence of a Britain-centered mode in the Iron Age for a modern British sample—that reflect temporal shifts in genetic similarity between ancient and modern populations [3]. The interesting implication of this framing is that, in retrospect, the concept of “admixture” also takes on a different perspective. Not only is admixture conditional on the coordinate space choice (e.g., PCA spanned by continental references vs. references within-continent), but it is also dependent on the time period, which makes it a relativistic concept and measure, arguing against the classical presentist approach towards ancestry inference.

Several limitations follow directly from the framework and should guide interpretation. First, coordinate predictions are inherently conditional on the embedding space: the number and type of sampled populations represented in creating it will shift the space’s axes. Continuous coordinates reduce coercive categorization but do not eliminate the social and ethical risks of misinterpretations of genetic ancestry. Second, the resolution of both coordinates and breakpoint labels is tied to the windowing scheme, motivating the use of robust pooling and masked losses, which can be handled with caution. Third, geographic representations, in particular, add interpretability, but also introduce assumptions— especially for ancient samples, where the excavation site may not coincide with birthplace and where sampling coverage is more sparse. These caveats do not undermine the utility of coordinate-based LAI; rather, they emphasize that PCLAI’s outputs are only as good as its inputs, which will improve with increased sequencing of modern and ancient human populations.

Overall, PCLAI opens a new perspective for defining population descriptors, most importantly, defining “ancestry” as a population descriptor that has both discrete and continuous components, with the continuous component being multidimensional—as we have shown, and varying through space and time.

## Software Availability

The PCLAI source code and additional resources are publicly available on GitHub (https://github.com/AI-sandbox/pclai). PCLAI uses the VCF reader from the *snputils* library [76].

## Acknowledgments

We would like thank two former students: Richa Rastogi for demonstrating the coordinate-based LAI concept using an LSTM and transformers, results which were presented as a poster at the *Learning Meaningful Representations of Life* workshop at Neurips, Dec. 2020, “Ge2Net: Enabling genetic medicine for diverse populations by inferring geographic ancestry along the genome”, and Arvind Kumar for initial exploration of the coordinate-based concept using XGBoost [77], presented as a poster in the *AI for Affordable Healthcare* (AI4AH) workshop at ICLR, April 2020. In addition, we thank Peter Gerlach for manuscript feedback, and the UCSC Genomics Institute for providing computational resources and support during this project.

## Supplemental Material

**Table 1.**
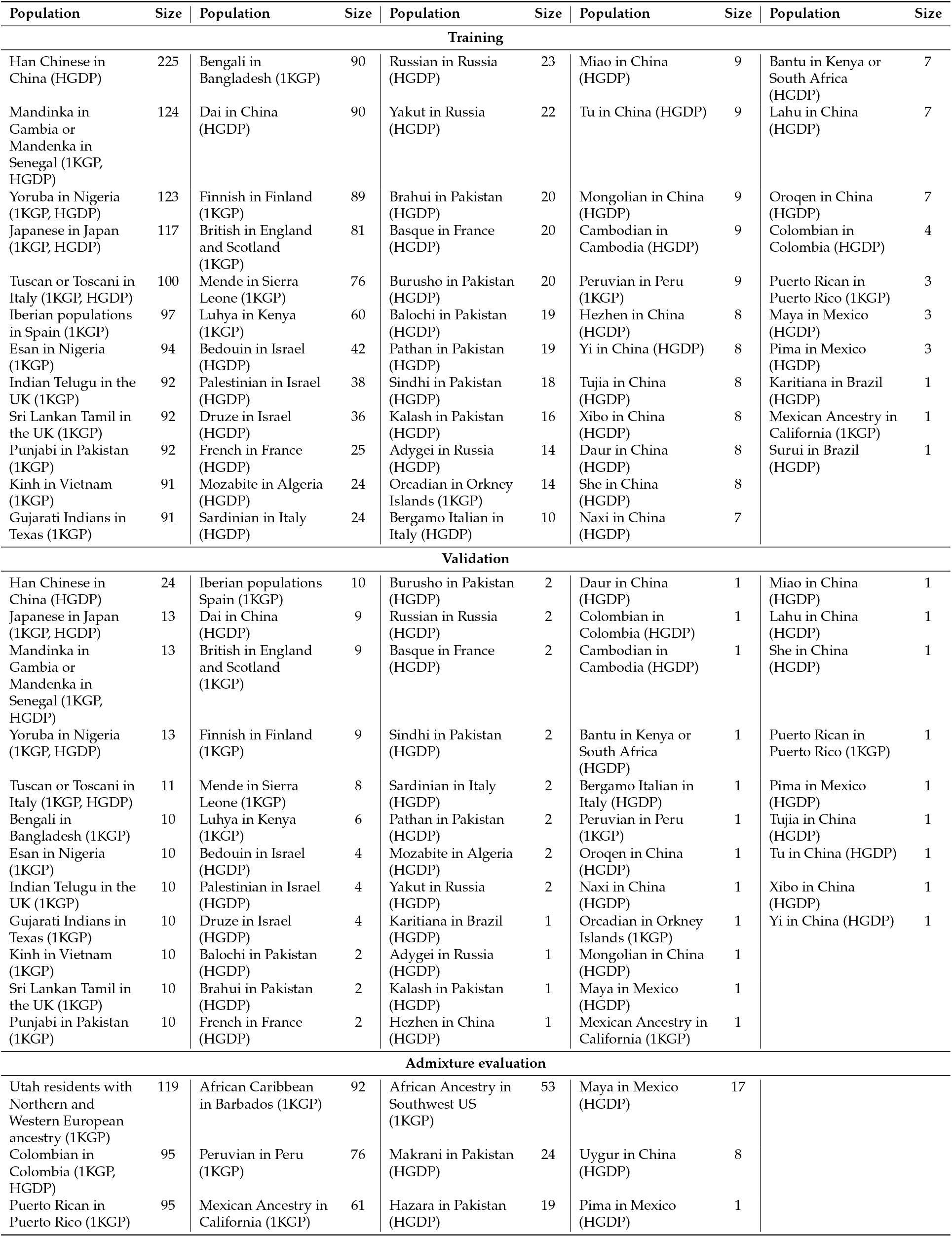
Human genomes dataset (from 1KGP [45] and HGDP [53])

**Table 2.** South Asian genomes dataset (from GenomeAsia 100K [49, 50])

| Population | Size | Population | Size | Population | Size | Population | Size | Population | Size |
| --- | --- | --- | --- | --- | --- | --- | --- | --- | --- |
| **Training** |  |  |  |  |  |  |  |  |  |
| Pakistani in Pakistan | 35 | Kalash | 18 | Lambada in Nilgiri Hills | 14 | Kota in Nilgiri Hills | 9 | Irulan | 3 |
| Bangladeshi | 35 | Gujjar | 18 | Halba in Chattisgarh | 13 | Dhurwa in Jharkand | 9 | Munda in Bihar | 2 |
| Punjabi in Pakistan | 35 | Toda in Nilgiri Hills | 18 | Agharia in Orissa | 12 | Chamar in Punjab and Haryana | 8 | Manipuri in Manipur | 2 |
| Pathan in Pakistan | 33 | Parsi | 17 | Tanti in West Bengal | 12 | Jamatia in Tripura | 8 | Kapu | 1 |
| Balochi | 35 | Bhanusali | 17 | Punjabi Brahmin | 12 | Koya Dora in Andhra Pradesh | 8 | Khonda Dora in Andhra Pradesh | 1 |
| Brahui in Pakistan | 33 | Oraon in Bihar | 16 | Abujmaria in Chattisgarh | 11 | Patel | 7 | Bagdi in West Bengal | 1 |
| Sindhi in Pakistan | 32 | Khatiri in Amritsar | 16 | Dorla in Chattisgarh | 11 | Chenchu in Andhra Pradesh | 7 | Maratha | 1 |
| Burusho in Pakistan | 29 | Birhor in Jharkhand | 16 | Kamar in Chattisgarh | 11 | Maharashtra Brahmin | 6 | Mala | 1 |
| Mog in Tripura | 22 | Balmiki in Haryana | 15 | Ramgharia in Punjab | 10 | West Bengal Brahmin in West Bengal | 6 | Madiga | 1 |
| Ahir in Haryana | 18 | Lodha in West Bengal | 15 | Muria in Chattisgarh | 10 | Santhal | 6 | Relli | 1 |
| Mahar in Punjab and Haryana | 18 | Konda Reddy in Andhra Pradesh | 15 | Paniya in Kerala | 10 | Sourastra Brahmin in Maharashtra | 5 | Yadava | 1 |
| Rajput | 18 | Saryupari Brahmin in Chharrigarh | 14 | Hill Korwa in Jharkhand | 10 | Iyengar in Tamilnadu | 5 |  |  |
| Meo in Haryana | 18 | Iyer in Tamilnadu | 14 | Toto in West Bengal | 9 | Bison Horn Maria in Chattisgarh | 3 |  |  |
| **Validation** |  |  |  |  |  |  |  |  |  |
| Pakistani in Pakistan | 35 | Mog in Tripura | 2 | Bhanusali | 1 | Maharashtra Brahmin | 1 | Patel | 1 |
| Bangladeshi | 35 | Chenchu in Andhra Pradesh | 1 | Bison Horn Maria in Chattisgarh | 1 | Munda in Bihar | 1 | Santhal | 1 |
| Punjabi in Pakistan | 12 | Dhurwa in Jharkhand | 1 | Abujmaria in Chattisgarh | 1 | Madiga | 1 | Saryupari Brahmin in Chattisgarh | 1 |
| Pathan in Pakistan | 4 | Dorla in Chattisgarh | 1 | Konda Reddy in Andhra Pradesh | 1 | Kota in Nilgiri Hills | 1 | Sourastra Brahmin in Maharashtra | 1 |
| Brahui in Pakistan | 3 | Halba in Chattisgarh | 1 | Khatiri in Amritsar | 1 | Lodha in West Bengal | 1 | Tanti in West Bengal | 1 |
| Sindhi from Pakistan | 3 | Hill Korwa in Jharkhand | 1 | Koya Dora in Andhra Pradesh | 1 | Lambada in Nilgiri Hills | 1 | Toda in Nilgiri Hills | 1 |
| Burusho in Pakistan | 3 | Irulan | 1 | Kapu | 1 | Parsi | 1 | Toto in West Bengal | 1 |
| Balochi | 3 | Iyengar in Tamilnadu | 1 | Kamar in Chattisgarh | 1 | Paniya in Kerala | 1 | West Bengal Brahmin in West Bengal | 1 |
| Rajput | 2 | Chamar in Punjab and Haryana | 1 | Kalash | 1 | Oraon in Bihar | 1 | Yadava | 1 |
| Gujjar | 2 | Balmiki from Haryana | 1 | Jamatia in Tripura | 1 | Muria from Chattisgarh | 1 |  |  |
| Meo in Haryana | 2 | Agharia in Odisha | 1 | Iyer in Tamilnadu | 1 | Punjabi Brahmin | 1 |  |  |
| Mahar in Punjab and Haryana | 2 | Bagdi in West Bengal | 1 | Manipuri in Manipur | 1 | Ramgharia in Punjab | 1 |  |  |
| Ahir in Haryana | 2 | Birhor in Jharkhand | 1 | Mala | 1 | Relli | 1 |  |  |
| **Admixture evaluation** |  |  |  |  |  |  |  |  |  |
| Birbhum | 10 | Singapore South Indian | 10 | Gaud in Odisha | 5 |  |  |  |  |
| Birbhum General | 10 | Scheduled Caste in Gujarat | 10 | Nababuddha in Maharashtra | 4 |  |  |  |  |
| Birbhum OBC | 10 | Urban Coimbatore | 10 | Tamil | 3 |  |  |  |  |
| Birbhum SC | 10 | Sikh in Punjab | 6 | Brahmin | 2 |  |  |  |  |
| Birbhum ST | 10 | Urban Chennai | 5 | Telugu | 1 |  |  |  |  |

**Table 3.**
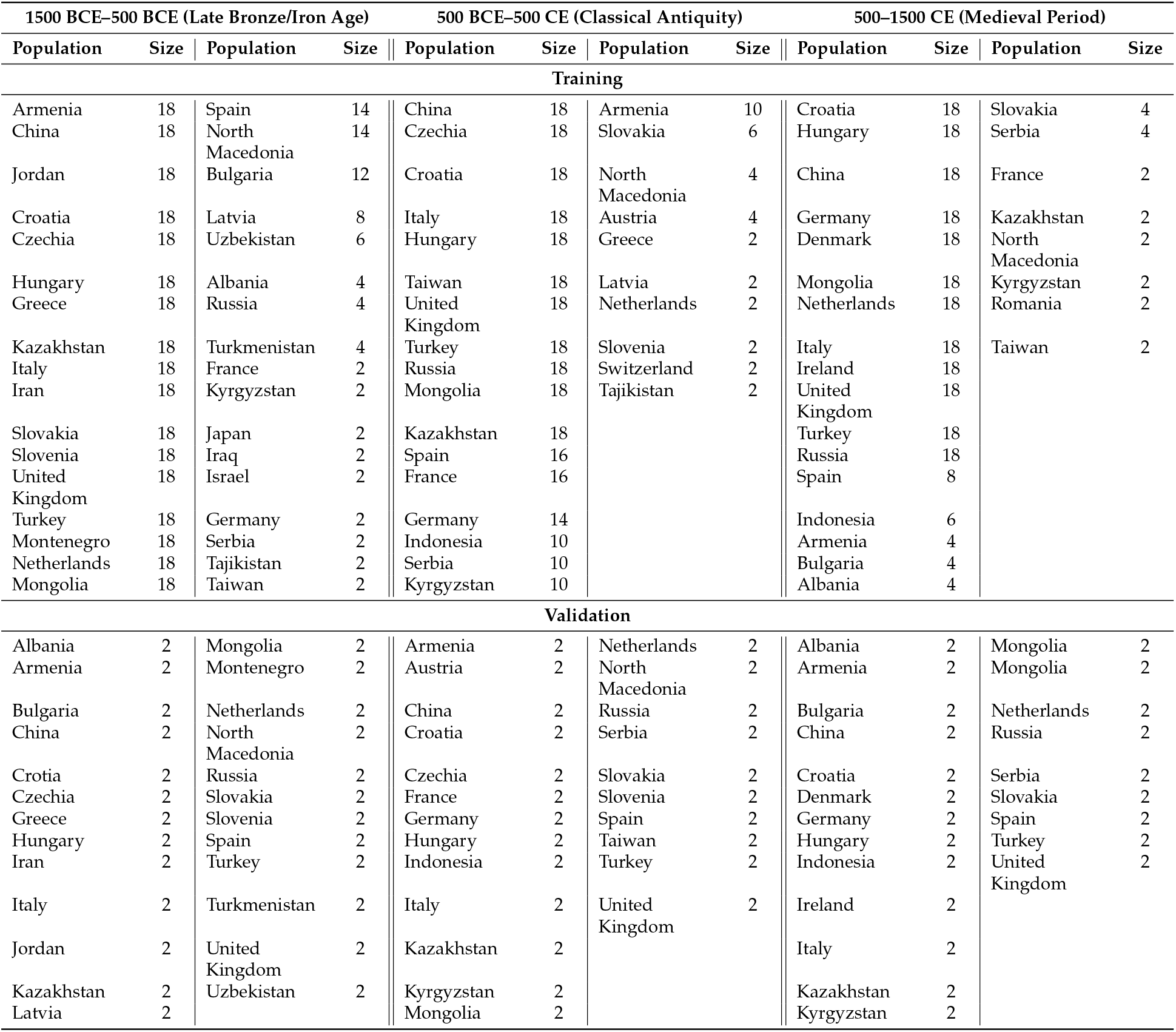
Ancient genomes dataset (from AADR [52])

**Figure S1.**
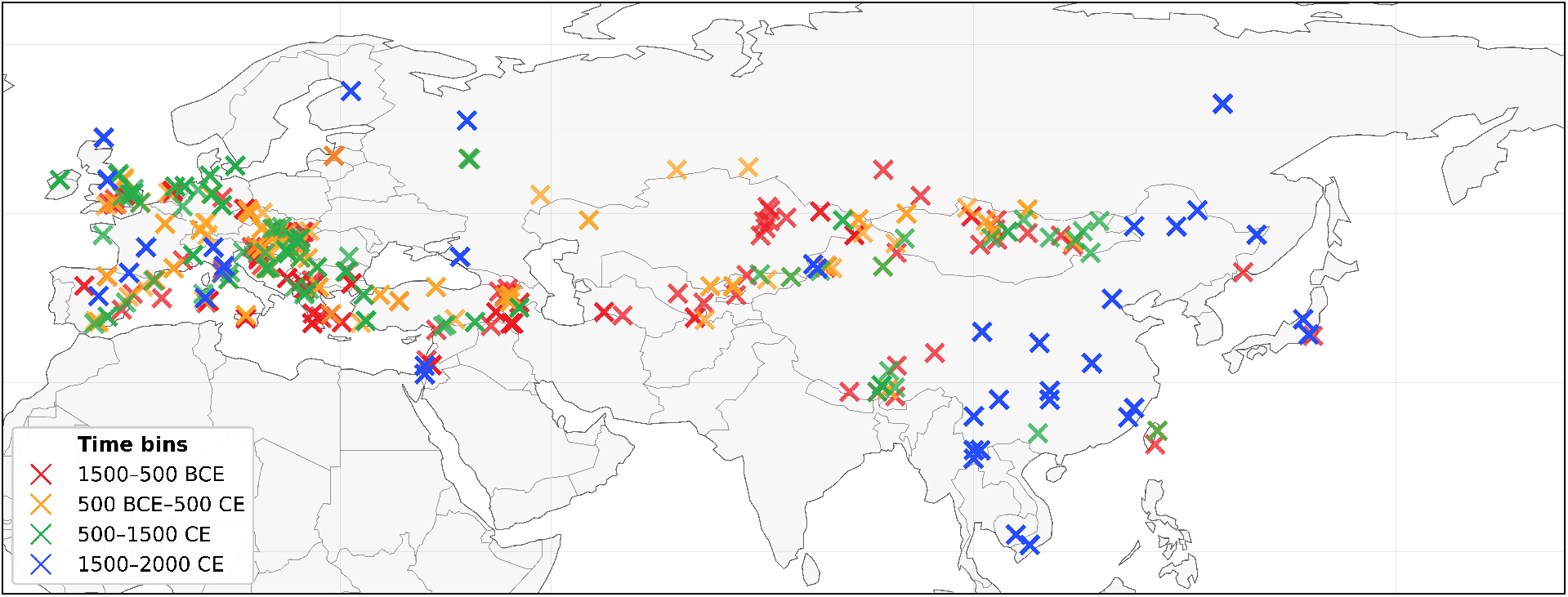
Geographic coverage of AADR ancient samples [52], complemented with modern samples from 1000 Genomes [45] and HGDP [53]; colored by time period.

## Supplemental Method S1: Hyperparameter tuning

To determine appropriate values for the Chamfer loss weight *γ*_geom_, the BCE changepoint loss weight *γ*_bp_, and the dropout probabilities in the transformer and MLP blocks, we performed a grid-search hyperparameter sweep. Each hyperparameter configuration was trained using five independent random seeds to account for stochastic variation in the model weight initialization and optimization with early stopping the same as in the default settings in the main text and the remaining architecture and loss weights held fixed during the sweep. Model quality was assessed using the unscaled raw ℒ_coord_ + ℒ_geom_ + ℒ_bp_ computed over 48, 000 held-out samples. Pairwise differences between hyperparameter settings were evaluated with Wilcoxon rank-sum tests, using a significance threshold of *α* = 0.1 and BH correction, appropriate for exploratory hyperparameter tuning, although some comparisons were significant at the *α* = 0.05 level as well. Our results in this section are *suggestive*, as with five runs we have low statistical power. Boxplots and statistical comparisons are shown in **Supplemental Figure S2**.

**Figure S2.**
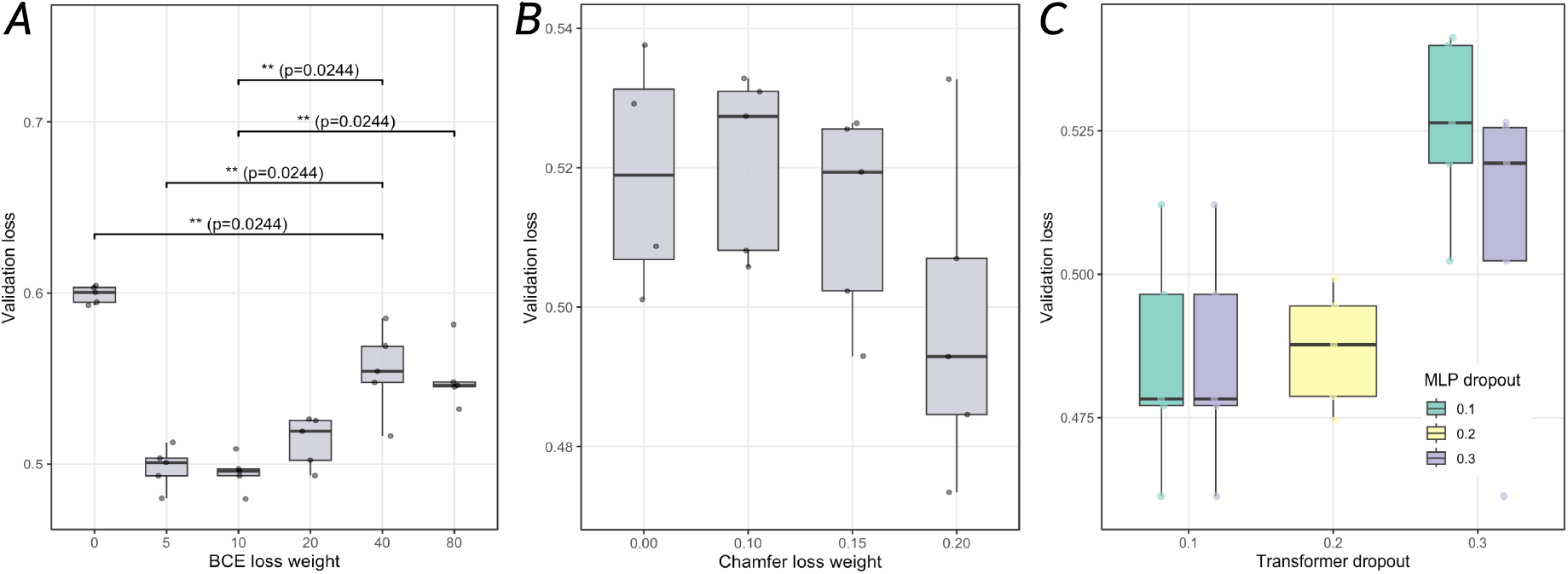
Hyperparameter tuning results. Validation loss was computed on 48, 000 held-out samples; *A*. Boxplots showing the distribution of validation loss across five training runs for each tested value of the Chamfer loss weight {0, 0.1, 0.15, 0.2 }; *B*. Boxplots showing the distribution of validation loss across five training runs for each tested value of the BCE loss weight {0, 5, 10, 20 }; *C*. Boxplots showing the distribution of combinations of MLP-transformer dropouts.

We evaluated the Chamfer loss weight *γ*_geom_ ∈ {0, 0.1, 0.15, 0.2}, and the results showed that there were no statistically significant differences between any pair of weights (all *p* ≥ 0.27), and all configurations produced very similar raw validation losses. Nevertheless, *γ*_geom_ = 0.2 consistently achieved the lowest median validation loss and the best individual runs, so we used *γ*_geom_ = 0.2 in subsequent experiments. For the BCE changepoint loss weight *γ*_bp_ ∈ {0, 5, 10, 20, 40, 80}, the results showed that introducing any non-zero BCE weight substantially reduced the validation loss compared to *γ*_bp_ = 0, and Wilcoxon tests indicated a significant improvement for all non-zero settings. There was a significant improvement at *γ*_bp_ = 5 compared to 0 at *p* = 0.0122, and marginal improvements at *γ*_bp_ = 20 compared to 0 (*p* = 0.0814) and at *γ*_bp_ = 5 compared to 10 (*p* = 0.0947). Overall, the setting *γ*_bp_ = 20 produced noticeably higher losses, while 5 and 10 were consistently lower. In light of these results we chose a moderate value, *γ*_bp_ = 10, which achieved low validation loss while keeping the BCE term numerically well-behaved and avoiding very large loss weights. Finally, we explored regularization by jointly tuning the transformer and MLP dropout probabilities. We compared three combinations: transformer dropout 0.1 with MLP dropout 0.3, transformer dropout 0.2 with MLP dropout 0.2, and transformer dropout 0.3 with MLP dropout 0.1 (**Supplemental Figure S2.C**). All three settings produced comparable performance, but the configuration with lower transformer dropout and higher MLP dropout (transformer 0.1, MLP 0.3) yielded the lowest median validation loss and slightly reduced variability across seeds; we therefore adopted this combination for the final model.

## Supplemental Result S1: Migration exploration in the *Familinx* dataset

When we use PCLAI to infer geographic coordinates, we make assumptions about where people are born. Some genomic datasets do not record birthplace at all (e.g., GenomeAsia 100K [49]) or simply cannot (e.g., ancient genomes from AADR [52]), as in the latter we assume that the place of burial is not *too far* from the true birthplace. But this may not always hold. To assess how reasonable these assumptions are, we used demographic data from the *Familinx* dataset [78] to assess human migration patterns in the past half millennium. We understand this does not generalize across all time periods and regions, but it offers a useful, empirical baseline for the magnitude of recent migration, helping contextualize how much geographic labels may deviate from true birth origin.

**Figure S3.**
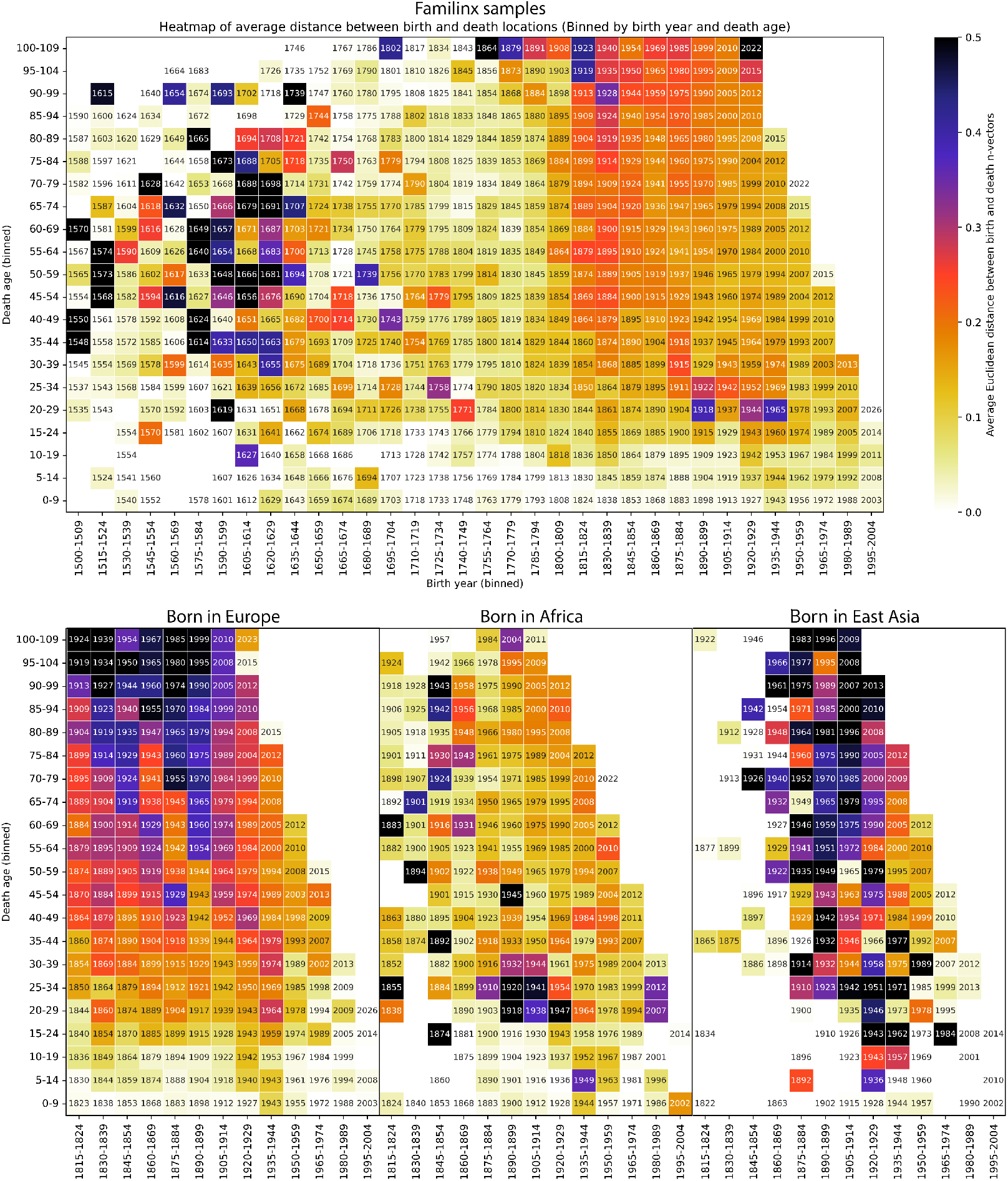
Heatmap of average distance between birth and death locations binned by birth year and death year in *Familinx* [78]. The year in each bin represents the average lifespan of binned individuals, i.e., the average year of death for those born in the year along the *x* axis who passed away at the binned death age along the *y* axis.

The *Familinx* subset provided to us contains 68, 586, 210 anonymized profiles of individuals scraped from Geni.com in 2015, of which 0.0039% (268, 290 profiles) have non-missing birth and death data if they were likely to be alive in 2015. The dataset also includes 51, 807, 144 parent-child relationships, with 0.434% (225, 341 edges) connecting complete profiles. Vital records, such as birth and death events, are significantly scarce, as confirmed by previous demographic studies such as [79], where it was reported that 60% of profiles are missing year of birth, and 42% of profiles with birth dates before 1942 are missing year of death. These issues are addressed by imputing missing birth and death years using recorded baptism and burial dates where available and necessary, and excluding profiles with implausible ages at death (>110 years). Additionally, the geographical information in the dataset is free-text, which is highly affected by plain orthographic errors, historical (defunct) names, and names in native languages, necessitating a substantial and time-consuming data cleaning process. Overall, the records exhibit significant geographic biases, with approximately 55% originating from Europe, 30% from North America, and the remainder distributed globally. In particular, most older records come from European and South Asian registers (dating from 1500 onwards), while the most recent record starts are predominantly from Pacific Island populations (dating from approximately 1780). Additionally, the *Familinx* dataset suffers from several biases, such as the under-representation of women, early deaths, marriages without children [80], and low- and middle-income countries, as well as a tendency towards *selective remembering* [79].

Following practices adopted in prior demographic studies [79, 81, 82, 83], we focused our analysis on individuals with non-missing birth and death years, as well as complete records of birth and death locations. Records predating 1600 are deemed unreliable and are generally excluded from the analysis— we still kept them in **Supplemental Figure S3** for transparency. Additionally, records after 1900 might include individuals still alive in 2015, potentially introducing ascertainment bias [81]. Even though there is significant variance in older records, interesting historical trends are still evident. We calculated the *n*-vectors from the latitude and longitude coordinates of birth and death locations, and computed the Euclidean distance between these *n*-vectors to quantify lifetime *migration* or, in other words, *displacement* for each individual. We then binned these values by birth year and age at death. One finding is that society began to be more panmictic in the 19th century, with individuals showing increased mobility from an early age. There are also high migration values during the 17th century, reflecting the first migration wave to the United States and, also, individuals who died during the world wars exhibited larger displacement values. This finding reinforces the results from [78], which showed that prior to the Industrial Revolution (before 1750), most marriages occurred between people born within 10 km from each other, while after the onset of the second Industrial Revolution (around 1870), marital distance increased rapidly, reaching approximately 100 km for those born in the 1950 cohort. At this micro-scale of human history, we can assume that in contemporary times there is a higher displacement throughout the lifetime.

## Supplemental Result S2: Additional PCLAI chromosome paintings

**Figure S4.**
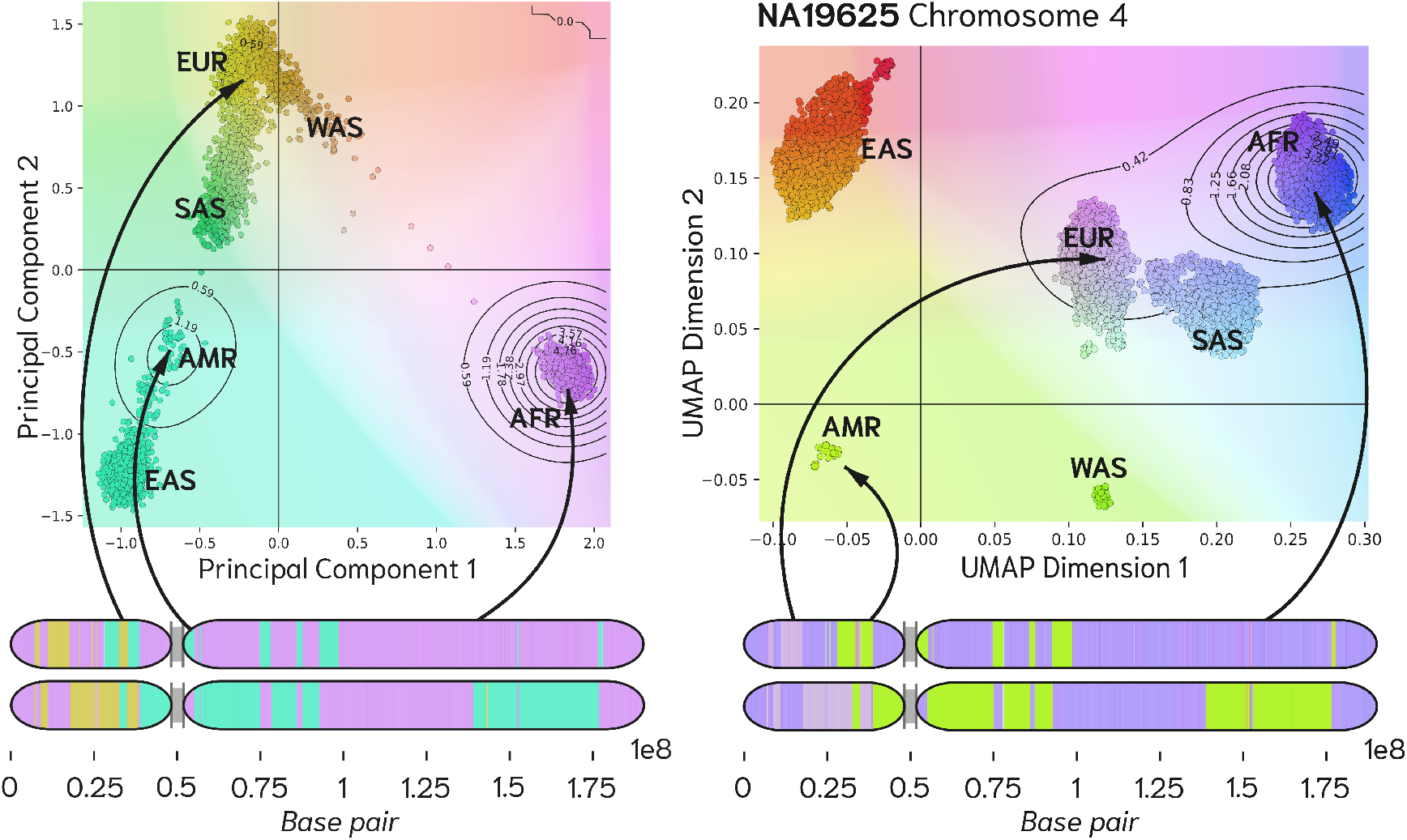
PCLAI Chromosome 4 paintings for a single African Ancestry in Southwest US individual in PCA (*left*) and UMAP (*right*) spaces. The bottom panels show haplotype PCLAI paintings of Chromosome 4 for NA19625 (African Ancestry in Southwest US). Colors along each chromosome correspond to the local coordinates predicted for 1,000-SNP windows. The top panels show founder haplotypes in the corresponding embedding spaces with continental groups labeled. Contour lines indicate the density of the NA19625 window-level chromosome-wide predictions projected into each space. Segments painted with colors that fall in the African-like cluster in PCA space tend to map to the African cluster in UMAP space, and similarly for other ancestries, illustrating that the two coordinate systems produce concordant ancestry attributions even though models had different training objectives and the color palettes differ.

In **Supplemental Figure S4** we illustrate that PCLAI produces visually consistent ancestry mosaics and breakpoint patterns under both linear (PCA) and non-Euclidean (UMAP) [43] embedding spaces. The bottom panels show Chromosome 4 paintings for NA19625 (African Ancestry in Southwest US), where each genomic window is mapped to coordinates in the reference embedding. The top panels overlay the corresponding predicted point cloud density contours on the reference space.

**Figure S5.**
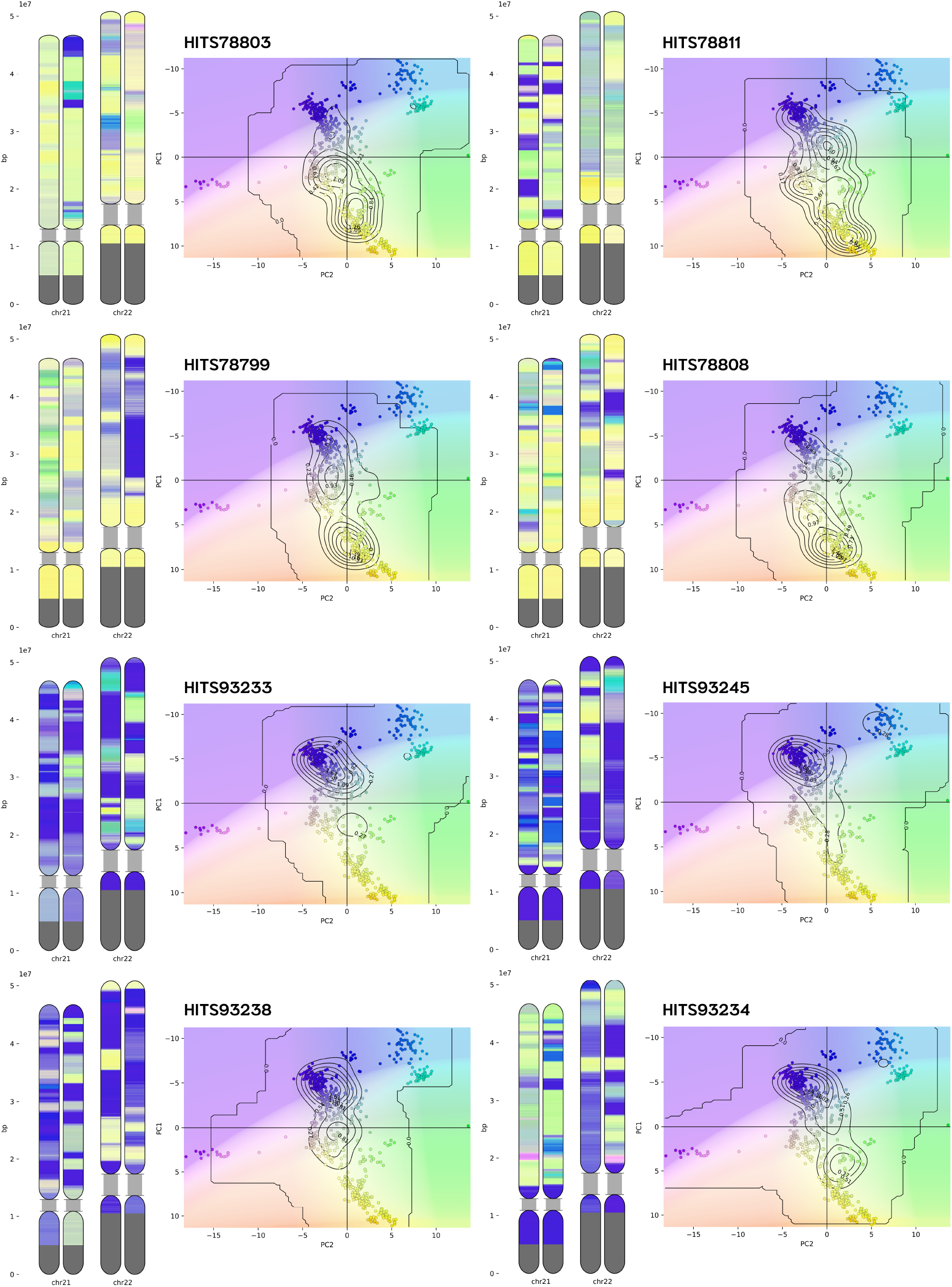
PCLAI chromosome paintings for South Asian individuals. *Left*: each row shows one GenomeAsia 100K [49] individual, with Chromosomes 21 and 22 paintings (*left*; two haplotypes per chromosome) and the corresponding distribution of inferred window-level coordinates in the South Asian PCA space (*right*; reference individuals as colored points, with density contours summarizing low-breakpoint-probability windows). The *top* four individuals are labeled as “Nababuddha in Maharashtra”, and the *bottom* four are “Scheduled Caste in Gujarat”.

## Supplemental Result S3: PCA of ancient genomes

We fit a PCA on our modern reference variation—4,091 European, West Asian, and East Asian samples from 1000 Genomes and HGDP (**Supplemental Table 1**) using max-heterozygosity 260,000 biallelic SNPs—and we projected our 1,945 AADR European, West Asian, and East Asian individuals (**Supplemental Table 3**) onto those axes. Because the PCA vectors are learned from modern genetic variation, projected ancient samples are read as positions relative to modern genetic drift directions. In **Supplemental Figure S6** we observe several patterns that align with historically documented mobility and admixture across Eurasia.

**Figure S6.**
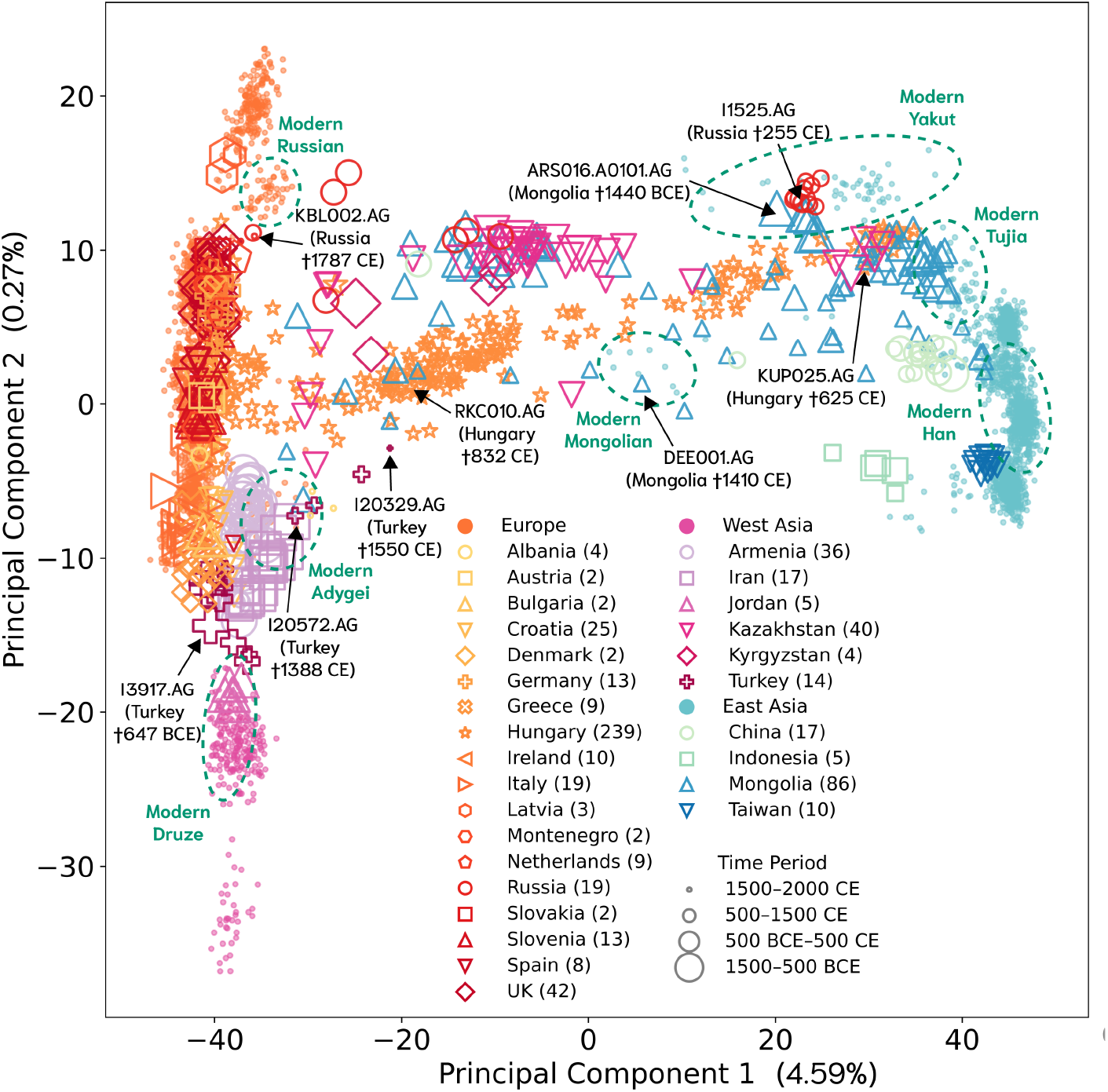
PCA of time-stratified ancient genomes projected onto modern Eurasian variation reveals trans-Eurasian clines and region-specific structure. Principal components are fit on present-day European, West Asian, and East Asian reference samples, and AADR [52] ancient individuals are projected onto the same axes. Point color indicates broad region (Europe—*Orange*, West Asia—*Pink*, East Asia— *Blue*), marker shape indicates country/region label (legend), and marker size encodes time bin (Late Bronze/Iron Age 1500–500 BCE, Classical Antiquity 500 BCE–500 CE, Medieval Period 500–1500 CE, Modern Era 1500–2000 CE). Dashed ellipses highlight clusters of present-day reference populations (e.g., Russian, Adygei, Druze, Mongolian, Yakut, Tujia, Han) for orientation. The percent variance explained by each axis is indicated.

1. *The Mongolian cline*: the most ancient samples found in present-day Mongolia (such as ARS016.A0101, † 1440 BCE) from the Late Bronze/Iron Age are genetically close to modern Yakut and Tujia, and to ancient samples found in Kazakhstan. Medieval samples from Mongolia (e.g., DEE001, †1410 CE) move toward the present-day Mongolian cluster.
2. *The Hungarian cline*: a larger set of medieval samples associated with the Carpathian Basin and Hungary are displaced from the core modern-day European cluster and extend toward Inner/East Asian variation. This pattern matches genetic evidence that nomadic confederations (e.g., Huns, Avars) entered the Carpathian Basin in successive waves, and that these waves included an immigrant component with an Inner Asian origin [84, 85] (e.g., KUP025, † 625 CE), so the PCA is reflecting this cline.
3. *The Turkish cline*: Iron Age samples found in Anatolia (e.g., I3917, † 647 BCE) have high affinity with modern West Asian/European samples, but in a later period the Classical Antiquity samples found in Turkey (e.g., I20572, † 1388 CE) show a consistent shift towards Caucasian populations such as present-day Adygei [86]. Later, during Medieval ages the samples move towards Central Asia (e.g., I20329, † 832 CE). This eastward pull is consistent with the historically documented Turkification of Anatolia following medieval Turkic expansions [87].
4. *Two ancient Russian clusters*: ancient Russian-associated individuals separate into two clusters in the projected space. Given Russia’s enormous geographic span and diverse ethnolinguistic histories, this split is plausibly capturing regional structure—one closer to modern Russian (e.g., KBL002, † 1787 CE) and the other to modern Yakut (e.g., † 255 CE). Notably, within each cluster we observe substantial overlap across time slices.

Overall, these PCA patterns reinforce the framing adopted by PCLAI: ancestry is a descriptor that changes across both space and time [6], shaped by genetic drift and admixture, so ancient individuals are not guaranteed to align ancestrally with modern individuals living at the same geographic locations [3].

## Notes

https://github.com/AI-sandbox/pclai

### Competing Interest Statement

The authors have declared no competing interest.

